# Thiooxazole Formation on a Nontypeable *Haemophilus influenzae* Virulence Factor Requires a Mixed-Valent Diiron Cofactor

**DOI:** 10.64898/2026.07.17.739211

**Authors:** Olivia M. Manley, Madeline B. Ho, Patrick M. McLean, Philip M. Palacios, Yisong Guo, Brian M. Hoffman, Amy C. Rosenzweig

## Abstract

The multinuclear nonheme iron-dependent oxidative enzyme (MNIO) family employs a multi-iron cofactor to catalyze a range of post-translational modifications (PTMs) in the biosynthesis of ribosomally synthesized, post-translationally modified peptide (RiPP) natural products. While significant progress has been made toward understanding the range of chemical transformations performed by MNIOs, the nature of the iron cofactor has only been investigated in one instance. Here, we examine the MNIO involved in oxazolin biosynthesis to gain further insight into the metallocofactors employed by this impressive family of enzymes. Oxazolin, a RiPP virulence factor from nontypeable *Haemophilus influenzae*, contains six copper-binding 5-thiooxazole groups installed by the MNIO HvfB. Weak interactions between HvfB and its required partner protein, HvfC, motivated genetic fusion of the two proteins, which yielded an effective mimic of the protein complex with high enzymatic activity. While HvfB binds up to three iron ions, concerted EPR, ENDOR, and Mössbauer spectroscopic characterization of the active protein reveals that accumulation of a mixed-valent diiron(II/III) cluster correlates with 5-thiooxazole product formation. This oxidation state is attained only in the presence of HvfC, revealing a new role for the partner protein in modulating the iron cofactor. Site-directed mutagenesis of metal-coordinating residues was used to probe the function of the third iron-binding site. This work clarifies the nature of the active iron cofactor for oxazolin maturation, providing a second example of a mixed-valent diiron oxidase in RiPP biosynthesis.

**TOC Graphic:** 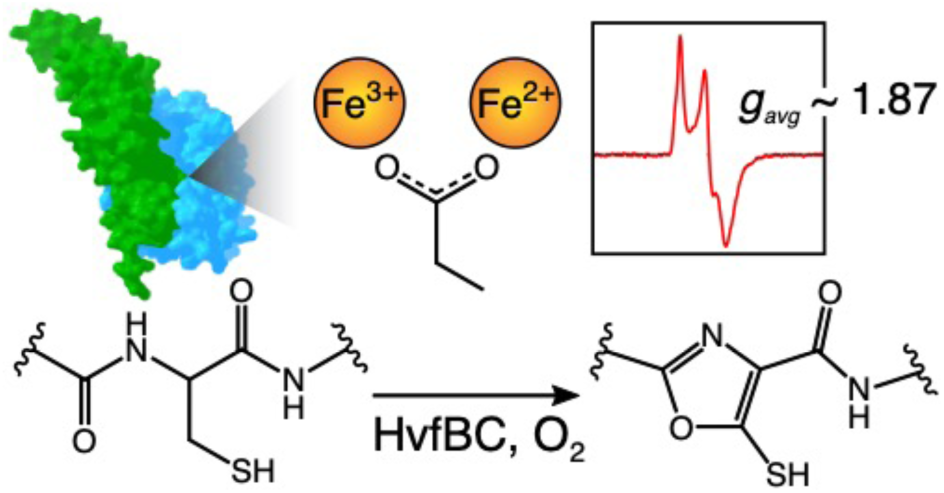

## INTRODUCTION

Multinuclear nonheme iron clusters are used as enzymatic cofactors to catalyze a broad array of chemical reactions in nature.^1–3^ These cofactors include carboxylate-bridged dinuclear clusters that are able to oxidize challenging substrates by coupling these reactions to the thermodynamically favorable reduction of dioxygen to water.^1^ A particularly well-studied example is soluble methane monooxygenase (sMMO), which uses a diferrous cluster to activate oxygen, traversing a series of established transient intermediate species to accomplish C–H cleavage and harness the ensuing oxygen rebound to convert methane to methanol.^4^ The catalytic mechanism of sMMO has served as a framework for understanding oxygen activation by diferrous clusters of numerous other diiron enzymes. However, a differing mechanistic framework is associated with the mixed-valent diiron oxidases/oxygenases (MVDOs), which as their name implies, use a mixed-valent Fe(II)/Fe(III) cluster for oxygen activation. As first exemplified by *myo*-inositol oxygenase (MIOX),^5, 6^ such mixed-valent clusters have since been identified as the active cofactors in fellow histidine-aspartate (HD)-domain MVDO proteins PhnZ and TmpB.^7–9^

The MVDO enzymes have recently expanded to include MbnB from *Methylosinus trichosporium* OB3b, the founding member of the multinuclear nonheme iron-dependent oxidative enzyme (MNIO) family. MbnB, in concert with its partner protein MbnC, uses a multinuclear iron cofactor to activate oxygen and convert cysteine residues to oxazolone and thioamide moieties for the biosynthesis of the copper-scavenging natural product methanobactin (Mbn) (Figure 1A).^10, 11^ Since its discovery, the iron cofactor in the MbnBC complex has been subjected to extensive spectroscopic interrogation. While Mössbauer spectroscopy and X-ray crystallography established that MbnBC can accommodate up to three iron ions in its active site,^10, 12, 13^ electron paramagnetic resonance (EPR) spectroscopy revealed an *S* = 1/2 signal attributed to a dinuclear Fe(II)/Fe(III) cluster (Figure 1B).^12^ The intensity of the mixed-valent EPR signal correlates with oxazolone/thioamide formation, suggesting that the diiron, rather than the triiron, form of the enzyme is the active species.^12^ Crystal structures obtained from low-iron preparations of MbnBC corroborate that the third Fe-binding site is unoccupied.^12, 13^ Furthermore, electron nuclear double resonance (ENDOR) studies demonstrated that the ferric iron of the Fe(II)/Fe(III) cluster is coordinated by two histidine ligands (site 1) and binds a substrate Cys thiolate, while the ferrous iron is coordinated by one histidine (site 2) and likely binds O_2_ for activation, providing a complete picture of the active site in the catalytically relevant oxidation state (Figure 1B).^14^

Since the discovery of MbnB, the MNIOs have emerged as a large family (>14,000 members) of iron-dependent metalloenzymes involved in the biosynthesis of ribosomally synthesized, post-translationally modified peptide natural products (RiPPs).^15^ RiPPs arise from genome-encoded precursor peptides that are produced by the ribosome and subsequently post-translationally modified by one or more biosynthetic enzymes to create the final product.^16, 17^ The MNIOs are a relatively recent addition to the classes of known RiPP-modifying enzymes, yet the family has already been shown to catalyze a wide variety of peptide post-translational modifications (PTMs), including macro- and heterocycle formation,^10, 18–21^ carbon excision,^22, 23^ hydroxylation,^24–27^ and amino acid cleavages.^28, 29^ Despite the large number of identified MNIOs, detailed structural and spectroscopic studies have only been reported for MbnB, leaving the generality of the Fe(II)/Fe(III) active cofactor unknown.

Recently, we reported that an MNIO from the human pathogen nontypeable *Haemophilus influenzae* produces a copper-binding RiPP virulence factor, termed oxazolin,^19^ that is important for infection by the bacterium.^30^ The MNIO, HvfB, is encoded by the *hvf* operon alongside the gene for a precursor peptide, HvfA, which contains an N-terminal secretion signal peptide and three AEGKCGEGKCG motifs toward the C-terminus (Figure 1C).^30^ The *hvf* operon also contains HvfC, which is a RiPP recognition element (RRE)^31^ domain-containing partner protein, and a yet uncharacterized DoxX-like protein, HvfX. Together with its required partner protein HvfC, HvfB oxidizes the six repeated C-terminal Cys residues of HvfA to copper-binding heterocycles. Although the HvfBC-conferred PTMs were originally described as oxazolone-coupled thioamides like those of Mbn, we^32^ and others^33^ have since reassigned the PTMs as 5-thiooxazole groups (Figure 1D). Initial characterization of the iron cofactor in HvfB by Mössbauer spectroscopy showed the presence of triferric and diferric species,^19^ but the oxidation state and nuclearity of the *catalytically active* cofactor were not established.

Herein, we report biochemical and spectroscopic characterization of HvfB and an HvfBC fusion protein that enables stoichiometric “binding” of the partner protein and exhibits high enzymatic activity. Through a combination of EPR, ENDOR, and Mössbauer spectroscopies, we dissected the nature of the iron cofactor in the fusion protein, revealing an *S* = 1/2 Fe(II)/Fe(III) signal that arises only in the presence of the partner protein, HvfC. Activity assays demonstrate that the accumulation of this mixed-valent cluster correlates with 5-thiooxazole formation, indicating that this species is the catalytically relevant form of the cofactor. While ENDOR spectroscopy suggests that the third Fe binding site adjacent to the Fe(II)/Fe(III) cluster is unpopulated, site-directed mutagenesis of putative third iron ligands demonstrate these residues serve an important functional role. Overall, these findings unify the cofactor requirements of MNIOs for Cys oxidations to 5-thiooxazole and oxazolone-coupled thioamides in RiPP biosynthesis.

**Figure 1.**
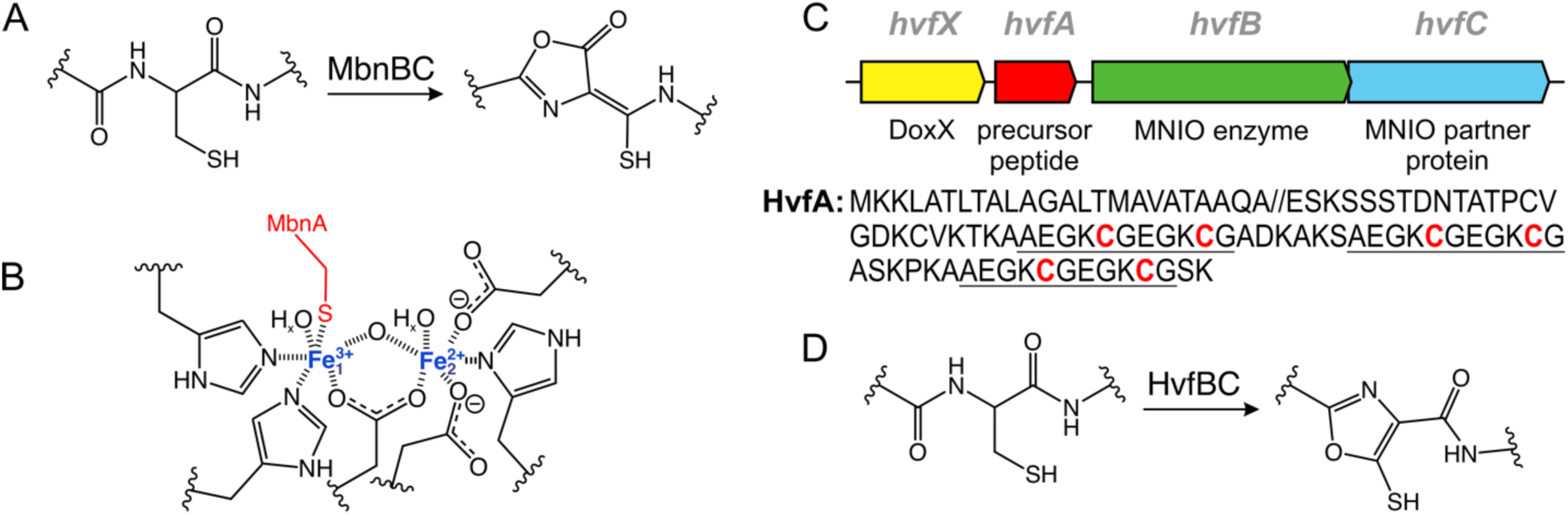
Overview of MNIOs MbnB and HvfB. (A) Modification of a cysteine residue to an oxazolone and thioamide for Mbn biosynthesis by MbnBC. (B) Schematic of the active MbnBC iron cofactor, with Fe site 1 and 2 labeled. (C) The *hvf* operon responsible for oxazolin biosynthesis. Gene names are listed above each gene with annotations below. The amino acid sequence of HvfA is shown with repeated motifs underlined and modified cysteine residues in red. The signal peptide cleavage site is denoted by “//”. (D) The conversion of a cysteine residue to a 5-thiooxazole moiety for oxazolin biosynthesis by HvfBC.

## EXPERIMENTAL SECTION

### Expression and Purification of *Hvf* Proteins

HvfA, HvfB, HvfC, and HvfBC_coexp_ were heterologously expressed in *Escherichia coli* NiCo21(DE3) cells and purified by Ni-affinity chromatography followed by size-exclusion chromatography as previously described.^19^ Minor differences from previous procedures were that cultures expressing HvfB were supplemented with 200 *µ*M ferrous ammonium sulfate at inoculation and again upon induction with 200 *µ*M IPTG. Additionally, for purification of HvfBC_coexp_, the protein was purified in one step by Ni-affinity chromatography to maintain as much complex as possible. Proteins were either dialyzed against 4 L of PBS at pH 7.8 or buffer exchanged via Amicon Ultra centrifugal filters (10 or 30K MWCO). Purified proteins were flash frozen in liquid N_2_ and stored at –80 °C until further use.

### Quantification of Iron Loading

Iron loading of HvfB was quantified by the ferrozine assay.^34^ Protein samples were treated with 1 M HCl to denature the protein and liberate the iron. The sample was then centrifuged at 21,000 × g for 10 min, and the supernatant was analyzed by ferrozine assay with comparison to ferric citrate standards ranging from 0 to 200 *µ*M.

### Expression and Purification of HvfBC_fusion_

The gene encoding HvfBC_fusion_ was synthesized by GenScript and subcloned into pET28a(+)-TEV at the NdeI and HindIII restriction sites for expression with an N-terminal, TEV-cleavable 6xHis-tag. The genetic fusion consists of *hvfB* immediately followed by *hvfC* with no linker between them (sequence available in Table S1). Expression and purification conditions for HvfBC_fusion_ were modified from those of HvfBC_coexp_ to enhance protein folding and activity. (Expression and purification of HvfB, HvfC, or HvfBC_coexp_ in the same manner as HvfBC_fusion_ did not have an effect except lowering yield, so the above conditions were used.) HvfBC_fusion_ was heterologously expressed in *E. coli* NiCo21(DE3) containing the pGro7 chaperone plasmid (Takara Bio) to overexpress GroES and GroEL. A culture in LB media containing 50 *µ*g/mL kanamycin sulfate and 30 *µ*g/mL chloramphenicol was grown overnight at 37 °C and used to inoculate 1 L of TB media supplemented with 50 *µ*g/mL kanamycin sulfate, 30 *µ*g/mL chloramphenicol, and 200 *µ*M ferrous ammonium sulfate in 2 L baffled flasks. Cultures were grown at 37 °C with shaking at 200 rpm until OD_600_ of 0.4-0.5, at which point chaperone expression was induced with 2 g/L L-arabinose, and the cultures were cooled to 18 °C. After 45 min, at an OD_600_ of 0.7-0.9, protein expression was induced with 50 *µ*M IPTG, and the cultures were supplemented with another 200 *µ*M ferrous ammonium sulfate. Variable iron loading was achieved by altering the amount of ferrous ammonium sulfate supplemented in the media, ranging from 0 to 200 *µ*M added at inoculation, induction, or both. For the lowest iron concentrations, expression was carried out in M9 minimal media to reduce metal availability further. After 20-22 h, cultures were harvested by centrifugation at 9,000 × g for 10 min, and the pellets stored at –20 °C until purification. For parity, purification by Ni-affinity chromatography was carried out in the same manner as described for HvfBC_coexp_ (ref^19^ and above). HvfBC_fusion_ was buffer exchanged and concentrated using Amicon Ultra centrifugal filters (50K MWCO), flash frozen, and stored at –80 °C until further use.

### In vitro Reactions of HvfBC

Reactions of HvfBC with HvfA were performed by combining 5 *µ*M enzyme [HvfB, HvfC, both (HvfBC_comb_), HvfBC_coexp_, or HvfBC_fusion_], 50 *µ*M HvfA, ∼1 mM O_2_ (from O_2_-saturated buffer ∼2 mM), and 1 mM dithiothreitol (DTT) at room temperature. DTT was added last to initiate the reaction. The reaction was then monitored on an Cary UV-visible spectrophotometer (Agilent). Spectra were collected every 30 s for 30 min. Reactions were stopped by the addition of 10x quench buffer (0.1 mM sodium citrate, 0.5 mM EDTA, pH 3.0, freshly prepared). The product was then analyzed using an Agilent 6230 LC-TOF mass spectrometer as previously described.^19^ All samples were treated with 1 mM DTT to reduce disulfide bonds prior to mass analysis, unless otherwise stated.

### Expression with ^57^Fe for Spectroscopic Studies

Due to low yields and poor enzymatic activity when expressed in M9 media, HvfBC_fusion_ was enriched with ^57^Fe in TB media. Culture flasks were soaked in dilute aqua regia prior to use. An LB starter culture of *E. coli* NiCo21(DE3) containing pET28a(+)-TEV-HvfBC_fusion_ and pGro7 was grown overnight with appropriate antibiotics as described above, and 10 mL of culture were used to inoculate 1 L TB media with antibiotics in 2-L acid-washed baffled flasks. ^57^Fe(s) was dissolved overnight in 1 M sulfuric acid (for a stock concentration of 100 mM ^57^Fe) inside a Coy anaerobic chamber the night before use. Solubilized ^57^Fe was removed from the anaerobic chamber and immediately added to the cultures to a final concentration of 200 *µ*M. The cultures were grown and induced with L-arabinose and IPTG as described for natural abundance preparations above. Upon induction, cultures were supplemented with another 200 *µ*M ^57^Fe added from the anaerobic stock solution. The remainder of the expression and purification were carried out as described above. Isotopic enrichment of ∼95% ^57^Fe was confirmed by inductively coupled plasma-mass spectrometry (ICP-MS, Thermo iCAP Q, Northwestern Quantitative Biological Imaging Facility) using samples prepared in the same manner as for ferrozine analysis.

### Sample Preparation for Spectroscopy

For EPR and Mössbauer spectroscopy, protein samples were made anaerobic using a Schlenk line, then transferred into a Coy vinyl anaerobic chamber where the rest of the manipulations were performed using anaerobic reagents. To prepare samples for X-band EPR, protein was concentrated to >0.3 mM using Amicon Ultra centrifugal filters before making it anaerobic and bringing it into the anaerobic chamber. For the as-isolated samples, no reductant was added. For the ascorbate titration, either 0.5, 1.0, 3.0, or 5.0 molar equivalents of ascorbate were added to the protein from a 10 mM stock solution. The fully reduced sample was prepared by addition of 20 *µ*M methyl viologen as a mediator and 5 molar equivalents of sodium dithionite, from a stock solution that was quantified using the absorbance at 315 nm (ε = 8 mM^−1^ cm^−1^). Final protein concentrations in all EPR samples were 0.3 mM unless otherwise stated. Samples were transferred to quartz EPR tubes and frozen in liquid nitrogen.

To prepare the HvfA substrate-bound EPR samples, HvfA was made anaerobic and brought into an anaerobic chamber. For samples made with HvfA containing disulfide bonds, HvfA was used as-is. For samples prepared with reduced HvfA, the peptide was treated with 10 mM DTT. A PD-10 desalting column was made anaerobic by bringing it into the anaerobic chamber, equilibrating with anaerobic buffer (PBS pH 7.8), passing through 3 mL of 0.5 mM sodium dithionite to scrub residual oxygen, and re-equilibrating with anaerobic buffer. At that point, the column was used to buffer exchange reduced HvfA and remove excess DTT. After buffer exchange, the concentration of reduced HvfA was determined by the BCA assay. HvfA was then added to HvfBC_fusion_ pre-treated with 1.5 molar equivalents of sodium ascorbate. Final concentrations of HvfA and HvfBC_fusion_ were 0.6 and 0.3 mM, respectively. The samples were transferred to quartz EPR tubes and frozen in liquid nitrogen.

For ENDOR spectroscopy, ^57^Fe-enriched protein at 2.1 mM was anaerobically combined with 1.5 molar equivalents of sodium ascorbate. The sample was transferred to a quartz capillary tube and frozen in liquid nitrogen. A control sample was prepared in the same manner with protein containing natural abundance Fe. For Mössbauer spectroscopy, ^57^Fe-enriched protein at 0.9 mM was anaerobically treated with 1.5 molar equivalents of sodium ascorbate. The protein was transferred to a Delrin Mössbauer cup and frozen in liquid nitrogen. A parallel X-band EPR sample was prepared to confirm accumulation of the mixed-valent state.

### EPR and ENDOR Data Collection

Continuous wave (CW) X-band EPR spectra were collected on a modified Bruker ESP-300 spectrometer with a custom program for data acquisition. A liquid helium flow Oxford Instruments ESR-900 cryostat was used for temperature control. Unless otherwise stated, EPR conditions were as follows: microwave frequency ∼9.3 GHz, temperature 12 K, 320 ms time constant, 10 G modulation, and 25 dB microwave attenuation. Spin quantitation of the X-band EPR samples was performed using a series of spin standards consisting of equine skeletal muscle myoglobin (Mb, Sigma) coordinated by azide (N_3_).^12^ These Mb-N_3_ standards were quantified using visible absorption of the heme Soret band before addition of azide. Pulsed Q-band EPR and ENDOR measurements were performed on a previously-described^35^ 35 GHz spectrometer with the SpinCore PulseBlaster ESR_PRO 400 MHz digital word generator and an Agilent Technologies Acquiries DP235 500 MS/s digitizer using SpecMan4EPR software.^36^ 35 GHz EPR and ENDOR studies were performed at 2K using a helium immersion dewar. ^57^Fe Davies ENDOR experiments were performed using the pulse sequence [π − T − π/2 − τ − π − τ – echo] with RF applied during interval T. The RF frequency was random-hopped within the spectrum range to minimize sweep artifacts.

### Mössbauer Data Collection

Mössbauer spectroscopy was carried out using two spectrometers with Janis Research SuperVaritemp dewars, which allow studies in applied magnetic fields up to 8 T at temperatures ranging from 1.5 to 200 K. The isomer shifts are reported relative to iron metal at 298 K. Spectral simulations were performed using the WMOSS software package and SpinCount software as previously described,^19^ and all spectral figures were prepared using SpinCount software.^37^

### Site-Directed Mutagenesis

Site-directed mutagenesis of putative iron-binding residues in HvfBC_fusion_ was performed by whole-plasmid amplification with iProof GC Master Mix (Bio-Rad) using semi-overlapping primers containing the desired mutation (Table S2). The PCR product was digested with DpnI (New England Biolabs) at 37 °C for 2 h to digest parent plasmid. The resulting product was transformed into *E. coli* DH5a cells, and resulting colonies were cultured, miniprepped, and screened for the desired mutation by sequencing (ACGT, Inc.). Successful constructs were transformed into *E. coli* NiCo21(DE3) and expressed and purified in the same manner as the wild-type (WT) proteins.

## RESULTS

### In vitro Reactivity of HvfBC

The modification of HvfA by HvfBC was initially established by heterologous coexpression of HvfB, HvfC, and HvfA in *E. coli*.^19^ To perform more detailed biochemical analysis, we developed an in vitro reaction using the absorbance at 302 nm to monitor reaction progress. Coexpressed HvfB and HvfC (HvfBC_coexp_), which was previously found to produce the most HvfBC heterodimer,^19^ was reacted with HvfA in the presence of DTT and O_2_-saturated buffer. A new absorbance peak at ∼302 nm appeared (Figure 2A), consistent with formation of the 5-thiooxazole product. Omission of any reaction component led to only modest changes in the absorbance features and no detectable modification by mass spectrometry (Figures 2B, S1-S2). Heterologously expressed HvfA purifies as a mixture of the full-length peptide (10,281.7 Da) and peptide lacking the N-terminal Sec signal sequence (7,981.9 Da).^19^ Intact protein MS of full-length HvfA following reaction with HvfBC_coexp_ showed an average mass of 10,261.4 Da, –20.3 Da from the mass of the unmodified peptide, consistent with the formation of five –4 Da 5-thiooxazoles on the six modifiable cysteine residues (Figure 2A inset, Table S3). Interestingly, HvfA lacking the signal sequence showed a mass loss of only –2.5 Da, consistent with less than one PTM per peptide, implying HvfBC preferentially modifies the full-length peptide. This is consistent with the Sec signal peptide functioning similarly to a RiPP “leader” peptide used for substrate recognition, much like in the biosynthesis of a fellow Sec signal-containing RiPP, bufferin.^38^

**Figure 2.**
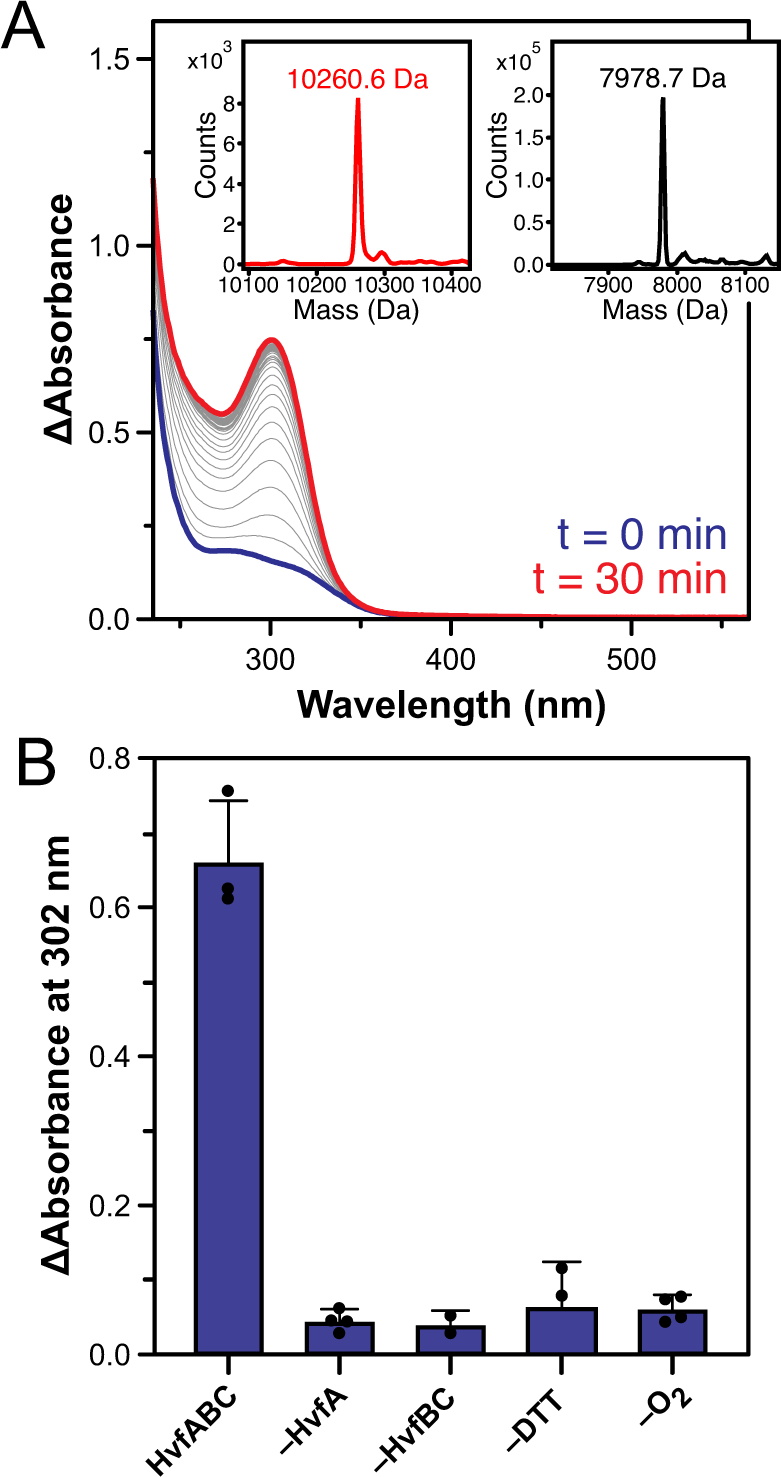
In vitro activity of HvfBCcoexp. (A) UV-visible spectra of the reaction progress, measured every 30 s (gray). The initial absorbance of unmodified HvfA alone is shown in blue, and the final absorbance of the reaction is shown in red. The spectrum of HvfBC was subtracted from the data for clarity. Reactions were performed using 50 *µ*M HvfA, 5 *µ*M HvfBCcoexp, 1 mM DTT, and ∼1 mM O2. Inset: The deconvoluted intact protein mass spectra of the full-length HvfA (red) and HvfA with the signal sequence cleaved (black) after reaction with HvfBC. (B) The increase in absorbance of the product chromophore at 302 nm for various reaction conditions. Spectral data are shown in Figure S1. Reported data are the average of at least three replicates with different protein preparations, and errors bars represent the standard deviation.

### Requirement for the Partner Protein

Most MNIOs characterized to date form a heterodimer with a partner protein that contains an RRE domain involved in substate recognition.^10, 13, 27, 39^ To probe the dependence of HvfB function on its partner protein HvfC, we investigated the reactivity of different preparations of the HvfBC complex. Omission of either HvfB or HvfC showed no product formation (Figures 3A, S3-S4, Table S4). Combining HvfB and HvfC (HvfBC_comb_) similarly showed no activity. These results are consistent with our previous finding that HvfBC heterodimer partially forms only when the proteins are coexpressed (i.e., HvfBC_coexp_ is a mixture of HvfBC heterodimer and HvfB and HvfC monomers, and HvfBC_comb_ consists of only monomers).^19^ To further assess the importance of the HvfBC heterodimer, we constructed a genetic fusion of HvfBC (HvfBC_fusion_), in which the C-terminus of HvfB is directly fused to the N-terminus of HvfC to ensure a 1:1 stoichiometry. HvfBC_fusion_ activity was first verified by in vivo coexpression with HvfA, which produced fully modified product (Figure S5). HvfBC_fusion_ was then expressed and purified in the same manner as HvfBC_coexp_, yielding samples with an average of 2.5 ± 0.2 mol. equiv. Fe per protein and a size consistent with the HvfBC dimer (Figure S6). In vitro activity assays with HvfBC_fusion_ show similar dependence on O_2_ and reductant (Figure S7-S8, Table S5), and importantly, HvfBC_fusion_ displays markedly higher product formation than HvfBC_coexp_ (Figure 3A-B). Considering the iron loading of HvfBC_fusion_ is roughly the same as HvfBC_coexp_, we attribute the higher activity to enhanced interaction between HvfB and HvfC in the fusion protein.

**Figure 3.**
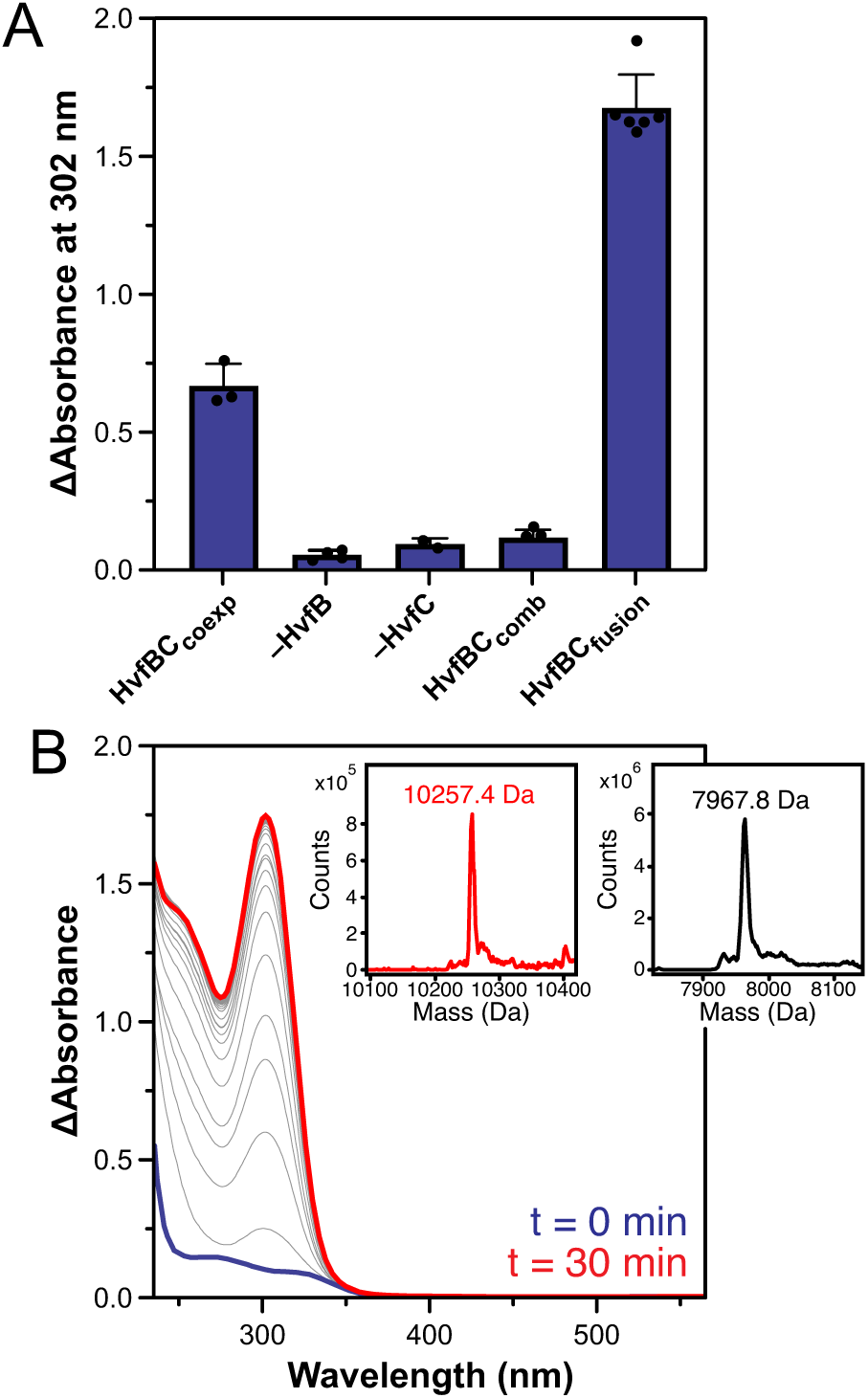
Dependence of HvfB activity on the formation of a complex with its partner protein, HvfC. (A) The increase in absorbance of the product chromophore at 302 nm of reactions containing HvfBCcoexp, omitting HvfB, omitting HvfC, combining HvfBC (HvfBCcomb), or using HvfBCfusion. Activity is reported as the increase in absorbance at 302 nm with the absorbance of unmodified HvfA subtracted. Reported data are the average of at least three replicates with different protein preparations, and errors bars represent the standard deviation. Corresponding spectral data are shown in Figs. 2A, S3, and panel B. (B) The timecourse of the reaction of HvfBCfusion with HvfA, with the initial spectrum of unmodified HvfA shown in blue and the final spectrum shown in red. Spectra were measured every 30 s (gray). The spectrum of HvfBCfusion was subtracted from the data for clarity. Reactions were performed using 50 *µ*M HvfA, 5 *µ*M enzyme (HvfB, HvfC, or HvfBCfusion), 1 mM DTT, and ∼1 mM O2. Inset: The deconvoluted intact protein mass spectra of the full-length HvfA (red) and HvfA with the signal sequence cleaved (black) after reaction with HvfBCfusion.

### Reductant and Iron Requirements

Because of its homogeneity and enhanced activity, we used HvfBC_fusion_ for further biochemical investigation. Since the reductant DTT was required for product formation (Figure S7), we speculated that it is serving to reduce disulfide bonds in the HvfA modifiable Cys residues and/or reduce the iron cofactor of HvfB for subsequent oxygen activation. HvfA and HvfBC_fusion_ were each deoxygenated and treated with reductant, desalted in an anaerobic glovebox, and then used for reactions. Combining DTT-treated HvfA with as-isolated HvfBC_fusion_ resulted in no product formation (Figure S9). Similarly, the reaction of DTT- or ascorbate-treated HvfBC_fusion_ with as-isolated HvfA also failed to generate product. Together, these data suggest that reductant is important both to prevent non-productive oxidation of the iron cofactor and to maintain the Cys thiols in the reduced state in order to be modified.

Previous work on MbnBC demonstrated that lower iron-loading resulted in higher enzymatic activity.^12^ To probe the effects of iron content, HvfBC_fusion_ was expressed in media supplemented with varying amounts of ferrous ammonium sulfate, resulting in different extents of iron loading in the purified protein. The activity of HvfBC_fusion_ was lower in preparations containing less than 2 molar equivalents of iron per protein, but similar in samples containing ∼2–3 molar equivalents of iron per protein, within typical variability across preparations (Figure S10). These results suggest that population of the third iron site is not required for activity, nor is it inhibitory.

### Mixed-Valent Fe(II)/Fe(III) State Correlates with HvfBC_fusion_ Activity

The X-band EPR spectrum of as-isolated HvfBC_fusion_ exhibits a rhombic signal with *g*-values of 1.96, 1.85, and 1.79, indicative of an *S* = 1/2 mixed-valent Fe(II)/Fe(III) cluster (Figure 4A) similar to that observed in MbnBC.^12^ A slight shoulder in the spectrum observed at *g* ∼ 1.98 may suggest some minor conformational heterogeneity. The *S* = 1/2 mixed-valent EPR signal is well-resolved at ∼12 K and diminishes in intensity at higher temperatures due to spin-lattice relaxation that reflects a relatively small-exchange coupling (*J)* between the two iron ions. This is consistent with other mixed-valent diiron centers, especially those with *μ*-hydroxo bridging ligands.^40^

Treatment of HvfBC_fusion_ with 1 molar equivalent of the reductant sodium ascorbate causes a modest increase in the mixed-valent signal (Figure 4A), with additional ascorbate failing to enhance the signal further (Figure S11). We note that ascorbate was not used in the above activity assays because its absorbance obscures the product peak. Instead, DTT was used, which similarly produces the mixed-valent signal (Figure S12). Spin quantitation of the *S* = 1/2 signal with maximum intensity demonstrates that it comprises ∼0.20 spins per protein (Figure S13), therefore ∼20% of the protein exists in the mixed-valent diiron form. The addition of either a strong reductant or oxidant diminishes the *S* = 1/2 species, corroborating its intermediate valence state (Figure 4A). Presumably, the site for the third Fe is unoccupied when the mixed-valent EPR signal is formed, as it is unlikely that a third iron ion could bind in such close proximity to the mixed-valent Fe(II)/Fe(III) cluster without participating in exchange coupling to the cofactor.

The spectrum of HvfBC_fusion_ displays an *S* = 5/2 signal at *g* = 4.3 in addition to the *S* = 1/2 signal from the mixed-valent cofactor (Figure 4A). Considering the iron stoichiometry of the sample (∼2.5 mol Fe/mol protein), we attribute this signal to a fraction of protein with a triferric cluster. Mössbauer spectroscopic data previously showed that HvfB, also isolated with 2.5 mol. equiv. of iron per protein, houses primarily an antiferromagnetically coupled triferric cofactor with a total spin of *S* = 5/2.^19^ Accordingly, the X-band EPR spectrum of HvfB itself displays a *g* = 4.3 signal that we attribute to the triferric cluster (Figure 4B, S14). Treatment with reductant fails to produce any of the *S* = 1/2 signal observed for HvfBC_fusion_ (Figures 4B, S14).

Ascorbate treatment of HvfBC_coexp_, which is made up of a mixture of HvfBC heterodimer and monomers, produces the mixed-valent signal, but less is accumulated compared to HvfBC_fusion_ (Figure 4B). The amount of mixed-valent species formed in HvfBC_coexp_ was somewhat variable between preparations but generally was at least 2-to-3-fold less than observed in the HvfBC_fusion_ samples (Figure 4B). The accumulation of the mixed-valent species thus appears to scale to the amount of HvfB bound to HvfC, indicating that the partner protein HvfC has a role in enabling the iron cluster to achieve the mixed-valent diiron state. Furthermore, the accumulation of the *S* = 1/2 signal also correlates with product formation across different preparations of the protein (HvfBC_fusion_ > HvfBC_coexp_ > HvfB, as shown above), positing this species as the catalytically relevant form of the cofactor.

**Figure 4.**
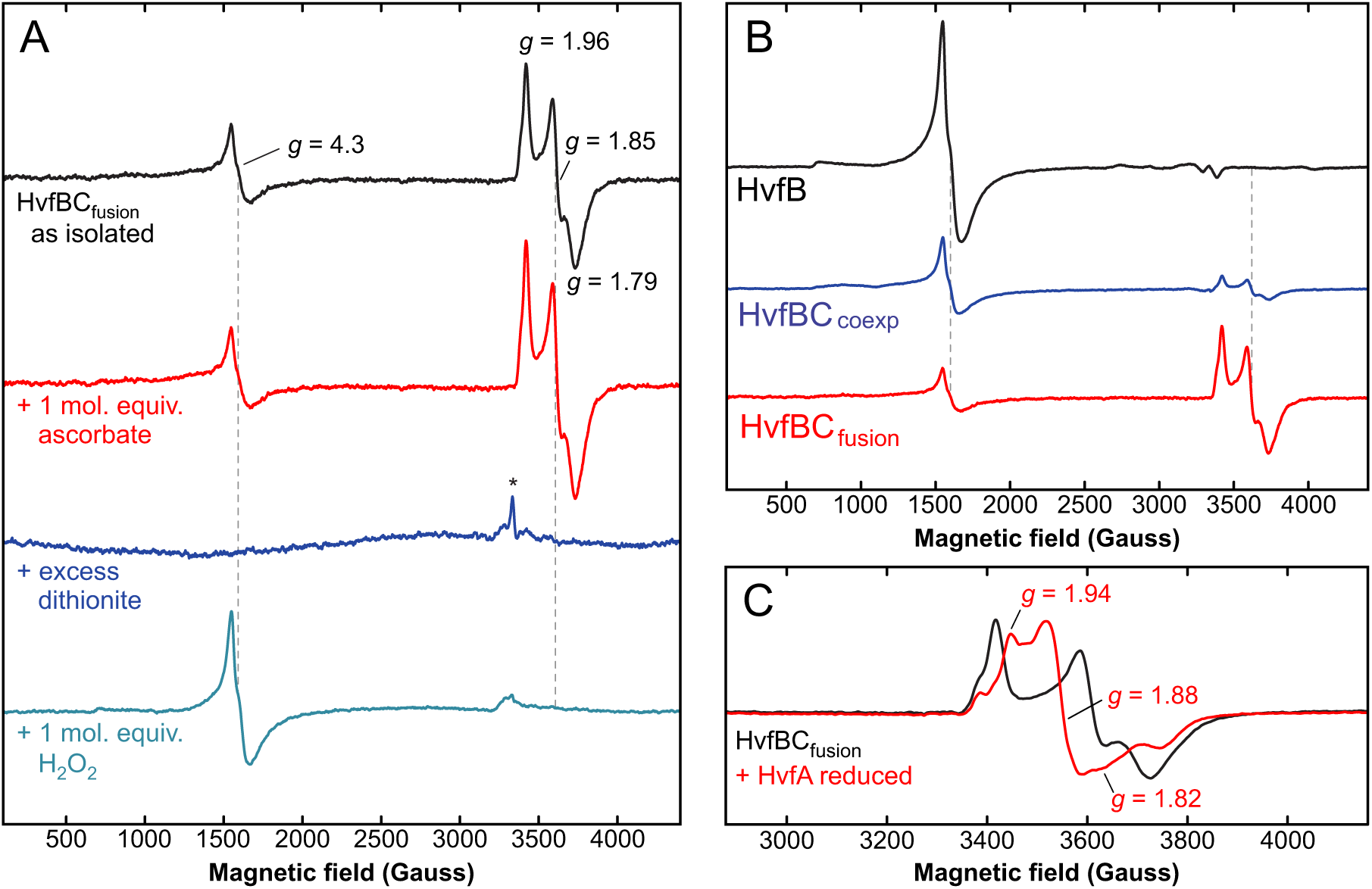
EPR spectroscopic evidence for a mixed-valent Fe(II)/Fe(III) cofactor in HvfBCfusion. (A) X-band EPR spectra of 0.3 mM HvfBCfusion as isolated (black) and treated with 1 mol. equiv. ascorbate (red), excess dithionite (blue), and 1 mol. equiv. hydrogen peroxide (teal). The asterisk represents some minor organic radical species, likely from the excess dithionite and methyl viologen. (B) X-band EPR spectra of 0.3 mM HvfB (black), HvfBCcoexp (blue), and HvfBCfusion (red) treated with 1.5 mol. equiv. ascorbate. (C) X-band EPR spectrum of 0.3 mM HvfBCfusion with the addition of 3 mol. equiv. of reduced HvfA (red), overlaid with the substrate-free spectrum (black). EPR conditions were microwave frequency of ∼9.3 GHz, temperature 12 K, 320 ms time constant, 10 G modulation, and 25 B microwave attenuation.

### HvfA Binding to HvfBC_fusion_

Coordination by substrate is known to perturb the EPR signal of mixed-valent diiron enzymes such as MbnB, MIOX, and others.^6–8, 12^ Because the modifiable Cys residues of HvfA are known to form disulfide bonds, HvfA was treated with DTT before combining with HvfBC_fusion_. The EPR spectrum of this combined sample indeed shows perturbation of the *S* = 1/2 signal (Figure 4C), suggesting that HvfA directly binds to the cluster. We note that the EPR spectrum generated from the addition of HvfA consists of multiple overlapping species. These features do not cleanly align with the EPR signal of HvfBC_fusion_ before addition of HvfA, and thus are not attributable to unbound cluster. Instead, the presence of multiple EPR signals can be attributed to conformational heterogeneity in the binding of HvfA substrate to the cluster. Considering the apparent majority species, the binding of HvfA results in a narrowing of the *g*-spread of the mixed-valent species to roughly *g* = [1.93, 1.88, 1.82] (Figure 4C). This *g*-tensor is consistent with sulfur ligation of one of the irons, like observed for MbnB.^12–14^ Addition of oxidized HvfA, in which the 6 modifiable Cys residues are in disulfide bonds, shows no change in the EPR signal (Figure S15), indicating that HvfBC is unreactive toward this form of the substrate and supporting the inference that HvfA binds to the cofactor via a cysteine thiolate.

### ENDOR Spectroscopy Corroborates Dinuclear Cluster

Q-band ^57^Fe ENDOR spectroscopy was used to further probe the *S* = 1/2 mixed-valent signal observed by EPR. For ^57^Fe-labeled HvfBC_fusion_ (2.3 mol Fe/mol protein), the spectrum shows two well-resolved hyperfine couplings centered at |A/2| = 18.5 MHz and 36.3 MHz, corresponding to the Fe(II) and Fe(III) sites of the mixed-valent cofactor, respectively (Figure 5A). We assign these signals based upon the well-known spin-projection coefficients for the ferric and ferrous sites of the *S* = 1/2 ground state of a spin-coupled diiron cluster, K_Fe(III)_ = +7/3 and K_Fe(II)_ = –4/3.^41^ Assuming the two irons have similar intrinsic ^57^Fe hyperfine couplings, the coupling with a larger magnitude is assigned to the ferric iron due to its larger spin-projection coefficient. Features at lower frequencies (< 15 MHz) observed in both the ^57^Fe-enriched and ^56^Fe natural abundance sample are assigned as ^14^N couplings from the coordinating histidine residues. The absence of a signal from a third iron in the ^57^Fe ENDOR spectra supports the conclusion, noted above, that the third iron-binding site is not occupied when the mixed-valent species is present. While a full 2D field-frequency ENDOR analysis to determine the complete hyperfine coupling tensors was not performed, we note that the ^57^Fe ENDOR in such multi-iron centers is typically dominated by the isotropic contribution, and the couplings observed at a single field are in broad agreement with those determined by Mössbauer spectroscopy (*vide infra*) and are also similar to those determined in the mixed-valent diiron cofactor of MIOX.^6^

**Figure 5.**
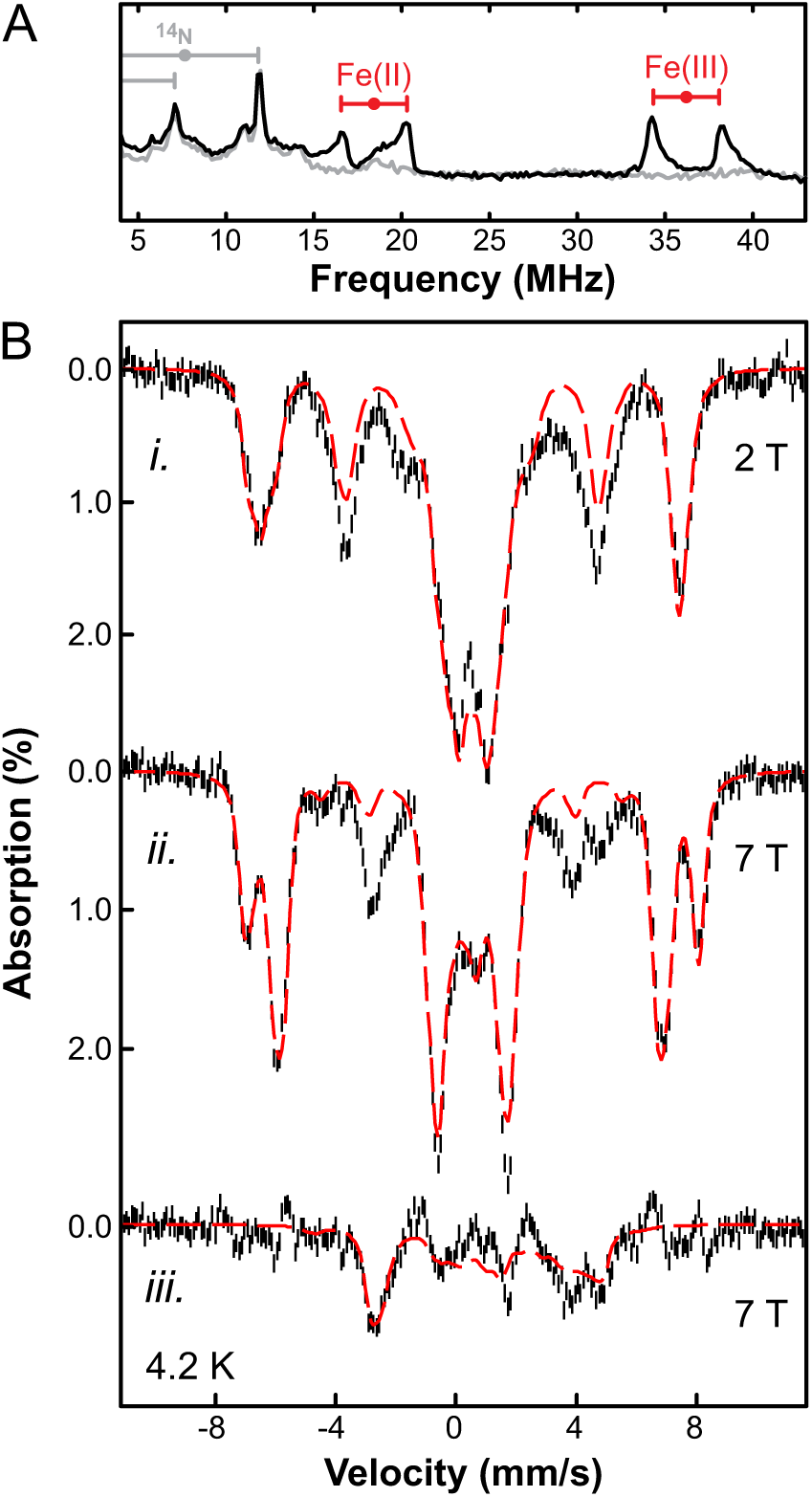
ENDOR and Mössbauer spectroscopy of ^57^Fe-labeled HvfBCfusion. (A) ^57^Fe Davies ENDOR spectra of the *S* = 1/2 mixed-valent species in isotopically-labeled ^57^Fe HvfBCfusion (black spectrum) and natural abundance HvfBCfusion (gray spectrum), collected at ∼13500 Gauss (∼*g*2). ^57^Fe couplings centered at A/2 and separated by 2\**ν*57Fe are denoted with red brackets. ^14^N couplings to Fe ligands are denoted with gray brackets. ENDOR conditions were ∼34.4 GHz, temperature 2 K, *τ* = 600 ns, TRF = 35 *μ*s, t90 = 40 ns, RF tail = 10 *μ*s, repetition time 50 ms. (B) 4.2 K Mössbauer spectra of ^57^Fe-enriched HvfBCfusion incubated with sodium ascorbate. The spectra were recorded under a 2 T (*i*) and 7 T (*ii*) applied magnetic field (black) and are shown overlaid with spectral simulations (red dashed lines), which are the sums of the simulations of triferric cofactor (∼55% of the total iron) and the diferric cofactor (∼25% of the total iron). (*iii*) The residual 7 T spectrum (∼20% of the total iron) was obtained by subtracting the 7 T simulation from the 7 T experimental spectrum shown in (*ii*); overlaid is a simulation as an *S* = 1/2 Fe(II)/Fe(III) mixed-valent cluster (red). Simulation parameters are listed in Table S6.

### Mössbauer Spectroscopy of HvfBC_fusion_

Since the mixed-valent diiron EPR signal accounts for only 20% of the protein, Mössbauer spectroscopy was used to gain further insight into the oxidation state of the remaining iron in the protein, which may be associated with the *g* = 4.3 EPR signal, representing the triferric cofactor, and/or may be EPR silent as an *S* = 0 antiferromagnetically coupled diiron species. The 4.2-K Mössbauer spectra of ^57^Fe-enriched HvfBC_fusion_ (2.3 Fe/protein) treated with 1.5 mol. equiv. of sodium ascorbate showed a dominant magnetic splitting feature, which can be simulated as 55% of the iron in the sample existing in the paramagnetic triferric state (Figure 5Bi-ii, Table S6), identical to the reported spectrum of primarily triferric HvfB.^19^

In addition, ∼25% of the spectral absorption is attributable to an *S* = 0 diferric state (Figure 6Bi-ii). Upon removing these two spectral components (the triferric and the diferric states) by subtracting their simulations from the spectral data, the remaining ∼20% absorption feature (Figure 6Biii) still exhibits a magnetic splitting pattern that can be reasonably well simulated as an *S* = 1/2 Fe(II)/Fe(III) mixed-valent cluster, with spectral parameters like that of the Fe(II)/Fe(III) mixed-valent cofactor found in MIOX (Figure 4Biii, Table S6).^6^ Thus, the Mössbauer analysis further corroborates the EPR/ENDOR results, supporting the existence of a dinuclear Fe(II)/Fe(III) mixed-valent cofactor in ∼20% of HvfBC_fusion_ upon ascorbate treatment, which we suggest is the active form of the cofactor based upon activity assays described above.

### Iron Site 3 Ligand Substitutions Hinder Activity

In MbnBC, the structurally characterized diiron cofactor is positioned in sites 1 and 2 (Figures 1B, S16),^12, 14^ and similarly, the crystal structure of an MNIO from *Histophilus somni* 129PT (PDB ID: 3BWW^42^) closely related to HvfB (72.4% identity, 83.8% similarity) also shows occupancy of a diiron cluster in sites 1 and 2 (Figure S16). AlphaFold models prepared with two iron ions per protein suggest that a diiron cluster may occupy the same position in HvfB, with inclusion of a third iron ion in the model needed for occupancy of site 3 (Figure 6A-B). To investigate the importance of the predicted site 3 ligands, these residues were individually replaced with alanine by site-directed mutagenesis. The HvfBC_fusion_ N175A and D242A variants each copurified with slightly lower iron content compared to the WT protein, while the Fe content of the H244A variant was similar to WT (Figure 6C). However, despite having bound iron, each variant was catalytically inactive (Figure 6D and Table S7).

X-band EPR spectra of ascorbate-treated N175A and D242A display significantly less intense *S* = 1/2 Fe(II)/Fe(III) signals than that of WT, even if accounting for decreased iron content, suggesting the majority of the iron is in an integer-spin state such as an *S* = 0 antiferromagnetically coupled diferric or diferrous cluster, though a small amount of the mixed-valent signal persists (Figure 6E). The H244A variant displays only a *g* = 4.3 signal consistent with high-spin *S* = 5/2 iron, reminiscent of the spectrum of triferric HvfB (Figure 4C). This observation is consistent with retention of the triferric cluster but an inability to achieve the active mixed-valent state of the cluster. More conservative Ser mutations also were made, with similar results to the Ala variants (Figure S17). Despite iron stoichiometry suggesting a diiron cluster may be present in each variant, the lack of activity and low intensity of the mixed-valent EPR signal suggest that these residues are important for forming or stabilizing the active iron oxidation state, despite iron not being bound in site 3.

**Figure 6.**
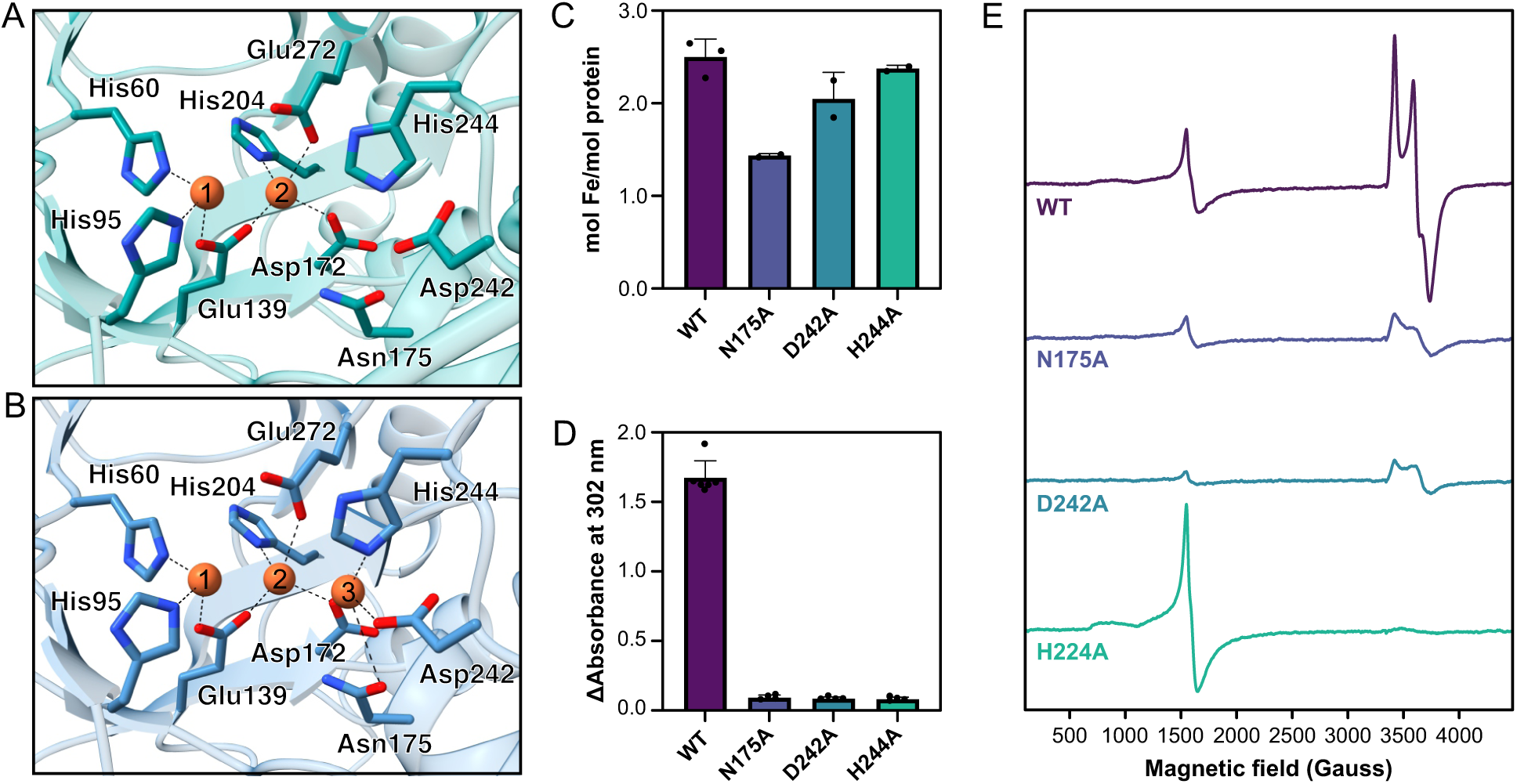
Roles of the predicted ligands to the third iron site. (A) AlphaFold model of HvfB with two or (B) three iron atoms. (C) Iron content of HvfBCfusion variants. (D) Activity of HvfBCfusion variants, measured by the increase in the product chromophore absorbance at 302 nm. Mass spectral analysis is shown in Table S7. (E) X-band EPR spectra of HvfB variants treated with 1 mol. equiv. of sodium ascorbate. EPR conditions were microwave frequency ∼9.3 GHz, temperature 12 K, 320 ms time constant, 10 G modulation, and 25 dB microwave attenuation.

## Discussion

With the reassignment by us^32^ and others^33^ of the six Cys modifications in oxazolin as 5-thiooxazoles rather than oxazolone and thioamide moieties, this work represents the initial investigation of the iron cofactor responsible for 5-thiooxazole formation by an MNIO. Spectroscopic data show that the majority of iron in HvfBC_fusion_ exists as high-spin ferric iron, seemingly as a triferric cluster like in HvfB.^19^ However, the presence of an *S* = 1/2 EPR signal arising from a mixed-valent diiron(II/III) cluster correlates with enzymatic activity, consistent with this being the active iron oxidation state. ^57^Fe ENDOR displays only two well-resolved couplings, consistent with the *S* = 1/2 EPR signal arising from two, not three, iron atoms. The Mössbauer spectra of HvfBC_fusion_ show a mixture of multinuclear iron species but corroborates that a proportion of the protein houses a mixed-valent Fe(II)/Fe(III) diiron cluster (∼20% of total iron). Together with the observation of similar activity for HvfBC_fusion_ samples containing 2-3 Fe ions per protein, these data indicate that occupation of Fe site 3 is not required for catalysis and that the enzymatic activity arises from the ∼20% of protein that exists in the mixed-valent Fe(II)/Fe(III) state, much like MbnBC,^12, 14^ thus unifying the active cofactor responsible for distinct Cys modifications by MNIOs.

The role of the proposed third iron-binding site in MNIOs has been a source of speculation since its detection.^10^ Many MNIOs copurify with nearly 3 molar equivalents of Fe per protein, and trinuclear iron clusters have been structurally verified in MbnB^12, 13^ and TglH^39^ and spectroscopically detected in MbnB^10^ and HvfB.^19^ The function of this third iron-binding site is especially puzzling since the mixed-valent *diiron* forms of MbnBC and, as shown here, HvfBC are reactive.^12, 14^ Correspondingly, crystal structures of MbnB with low iron loading^12^ and an HvfB homolog from *Histophilus somni* 129PT^42^ show preferential iron binding to sites 1 and 2, with site 3 having lower or no occupancy (Figure S16).

While our results indicate that occupancy of site 3 is not required for HvfBC enzymatic activity, site-directed mutagenesis of potential coordinating residues (N175, D242, and H244, Figure 6D-E, Figure S17) to either Ala or Ser completely abolished enzymatic activity and thwarted accumulation of the mixed-valent oxidation state, indicating that these residues are essential for HvfBC function. For comparison, substitution of the equivalent residues of MbnB with Ala or Ser shows diminished but detectable activity, slightly higher than the activity upon mutation of site 1 and 2 ligands.^10, 12, 13^ Our results are consistent with the interpretation that these site 3 residues function not in iron binding, but perhaps serve a structural or hydrogen-bonding role that is essential for enzymatic function.

This work also provides further insight into the role of the MNIO partner protein. Most MNIOs only function in complex with their partner proteins belonging to the PME (<u>P</u>artner protein of <u>M</u>NIO <u>E</u>nzyme) family, which consists of a DUF2063 domain fused to a full or partial RRE domain.^43^ Exceptions include MovX, which has a fused C-terminal RRE domain,^28^ and ChrH, which has an additional domain of unknown function.^15, 18^ MbnB can only be solubly expressed and purified from *E. coli* as a complex with its partner protein (MbnBC).^10^ Crystal structures of MbnBC with its precursor peptide MbnA implicate the importance of the PME1 RREs in coordinating substrate binding by interacting with the leader peptides.^13^ However, the inability to separate MbnB and MbnC precluded further investigation of the role of PME1 protein. Our comparison of HvfB, HvfBC_coexp_, and HvfBC_fusion_ demonstrates that in addition to the inferred role of HvfC in substrate binding, HvfC is required to achieve the active mixed-valent Fe(II)/Fe(III) state of the HvfB iron cofactor, thus expanding the functional understanding of the PME family.

The model of oxazolin maturation by HvfBC developed here begins with binding of the partner protein HvfC to HvfB, which enables formation of the catalytically relevant mixed-valent Fe(II)/Fe(III) oxidation state. HvfA binding results in coordination of the iron cofactor by the Cys thiol, presumably to the ferric iron, leaving the ferrous iron available for oxygen binding, as is suggested in MbnB (Figure 1B).^14^ While the mechanistic details following oxygen activation are yet unclear, it can be envisaged that the one-electron oxidative addition of O_2_ results in an Fe(III)/Fe(III)-superoxo intermediate, similar to those detected in other MVDOs like MIOX.^5^ Such a species may initiate 5-thiooxazole formation by C–H abstraction from the Cys Cβ. From there, several mechanistic possibilities for the formation of a 5-thiooxazole exist (Figure S18). Overall, the four-electron oxidation of Cys is balanced with the simultaneous reduction of O_2_ to water, resulting in regeneration of the active mixed-valent oxidation state at the end of the reaction, primed to bind and oxidize another Cys residue of HvfA until the six C-terminal Cys are all modified. Secretion into the periplasm and concurrent signal peptide cleavage result in mature oxazolin, which may either function in the periplasm or be exported by a yet unknown mechanism to carry out its functional role in copper binding. In summary, these findings support the requirement of a mixed-valent diiron cofactor by MNIOs performing Cys oxidations to both 5-thiooxazoles and oxazolone-coupled thioamides in RiPP biosynthesis.

## ASSOCIATED CONTENT

### Supporting Information

The Supporting Information contains supporting tables including DNA and amino acid sequences, mass spectrometry data, and Mössbauer simulation parameters, as well as supporting figures including mass spectra, UV-visible spectra, EPR and Mössbauer spectra, and a proposed catalytic mechanism (PDF).

### Notes

The authors declare no competing financial interests.

## ACKNOWLEDGEMENTS

This work was supported by NIH grants R35 GM118035 (A.C.R.), F32 AI176709 and K99 GM158961 (O.M.M.), R01 GM111097 (B.M.H.), and R01 GM125924 (Y.G.) and NSF CHE-2333907 (B.M.H.). We would like to acknowledge the Northwestern Integrated Molecular Structure Education and Research Center (IMSERC) for MS usage. IMSERC is supported by Northwestern University and the State of Illinois. We would also like to thank Rebecca Sponenburg for assistance with ICP-MS analysis, which was performed through the Northwestern Quantitative Biological Imaging Facility supported by the NIH grant S10OD020118.

## REFERENCES

(1) Jasniewski, A. J.; Que Jr, L. Dioxygen activation by nonheme diiron enzymes: diverse dioxygen adducts, high-valent intermediates, and related model complexes. Chemical Reviews 2018, 118 (5), 2554–2592. DOI: 10.1021/acs.chemrev.7b00457.

(2) Broadwater, J. A.; Haas, J. A.; Fox, B. G. The fundamental, versatile role of diiron enzymes in lipid metabolism. Lipid/Fett 1998, 100 (4-5), 103 – 113. DOI: 10.1002/(SICI)1521-4133(19985)100:4/5<103::AID-LIPI103>3.0.CO;2-4.

(3) Skirboll, S. S.; Ledinina, A. E.; Makris, T. M. Heme oxygenase-like dimetal oxidases and oxygenases. Biochemical Society Transactions 2026, 54 (3), 231–243. DOI: 10.1042/BST20253134.

(4) Banerjee, R.; Jones, J. C.; Lipscomb, J. D. Soluble Methane Monooxygenase. Annual Review of Biochemistry 2019, 88, 409–431. DOI: 10.1146/annurev-biochem-013118-111529.

(5) Xing, G.; Diao, Y.; Hoffart, L. M.; Barr, E. W.; Prabhu, K. S.; Arner, R. J.; Reddy, C. C.; Krebs, C.; Bollinger Jr, J. M. Evidence for C–H cleavage by an iron–superoxide complex in the glycol cleavage reaction catalyzed by *myo*-inositol oxygenase. Proceedings of the National Academy of Sciences 2006, 103 (16), 6130–6135. DOI: 10.1073/pnas.0508473103.

(6) Xing, G.; Hoffart, L. M.; Diao, Y.; Prabhu, K. S.; Arner, R. J.; Reddy, C. C.; Krebs, C.; Bollinger, J. M. A coupled dinuclear iron cluster that is perturbed by substrate binding in *myo*-inositol oxygenase. Biochemistry 2006, 45 (17), 5393–5401. DOI: 10.1021/bi0519607.

(7) Wörsdörfer, B.; Lingaraju, M.; Yennawar, N. H.; Boal, A. K.; Krebs, C.; Bollinger Jr, J. M.; Pandelia, M.-E. Organophosphonate-degrading PhnZ reveals an emerging family of HD domain mixed-valent diiron oxygenases. Proceedings of the National Academy of Sciences 2013, 110 (47), 18874–18879. DOI: 10.1073/pnas.1315927110.

(8) Rajakovich, L. J.; Pandelia, M.-E.; Mitchell, A. J.; Chang, W.-c.; Zhang, B.; Boal, A. K.; Krebs, C.; Bollinger Jr, J. M. A new microbial pathway for organophosphonate degradation catalyzed by two previously misannotated non-heme-iron oxygenases. Biochemistry 2019, 58 (12), 1627–1647. DOI: 10.1021/acs.biochem.9b00044.

(9) van Staalduinen, L. M.; McSorley, F. R.; Schiessl, K.; Séguin, J.; Wyatt, P. B.; Hammerschmidt, F.; Zechel, D. L.; Jia, Z. Crystal structure of PhnZ in complex with substrate reveals a di-iron oxygenase mechanism for catabolism of organophosphonates. Proceedings of the National Academy of Sciences 2014, 111 (14), 5171–5176. DOI: 10.1073/pnas.1320039111.

(10) Kenney, G. E.; Dassama, L. M.; Pandelia, M.-E.; Gizzi, A. S.; Martinie, R. J.; Gao, P.; DeHart, C. J.; Schachner, L. F.; Skinner, O. S.; Ro, S. Y.;, et al. The biosynthesis of methanobactin. Science 2018, 359 (6382), 1411–1416. DOI: 10.1126/science.aap9437.

(11) Reyes, R. M.; Rosenzweig, A. C. Methanobactins: Structures, Biosynthesis, and Microbial Diversity. Annual Review of Microbiology 2024, 78, 383–401. DOI: 10.1146/annurev-micro-041522-092911.

(12) Park, Y. J.; Jodts, R. J.; Slater, J. W.; Reyes, R. M.; Winton, V. J.; Montaser, R. A.; Thomas, P. M.; Dowdle, W. B.; Ruiz, A.; Kelleher, N. L.;, et al. A mixed-valent Fe(II)Fe(III) species converts cysteine to an oxazolone/thioamide pair in methanobactin biosynthesis. Proceedings of the National Academy of Sciences 2022, 119 (13), e2123566119. DOI: 10.1073/pnas.2123566119.

(13) Dou, C.; Long, Z.; Li, S.; Zhou, D.; Jin, Y.; Zhang, L.; Zhang, X.; Zheng, Y.; Li, L.; Zhu, X. Crystal structure and catalytic mechanism of the MbnBC holoenzyme required for methanobactin biosynthesis. Cell Research 2022, 32 (3), 302–314. DOI: 10.1038/s41422-022-00620-2.

(14) Jodts, R. J.; Ho, M. B.; Reyes, R. M.; Park, Y. J.; Doan, P. E.; Rosenzweig, A. C.; Hoffman, B. M. Initial steps in methanobactin biosynthesis: Substrate binding by the mixed-valent diiron enzyme MbnBC. Biochemistry 2024, 63 (9), 1170–1177. DOI: 10.1021/acs.biochem.4c00011.

(15) Chen, J. Y.; van der Donk, W. A. Multinuclear non-heme iron dependent oxidative enzymes (MNIOs) involved in unusual peptide modifications. Current Opinion in Chemical Biology 2024, 80, 102467. DOI: 10.1016/j.cbpa.2024.102467.

(16) Montalbán-López, M.; Scott, T. A.; Ramesh, S.; Rahman, I. R.; Van Heel, A. J.; Viel, J. H.; Bandarian, V.; Dittmann, E.; Genilloud, O.; Goto, Y.;, et al. New developments in RiPP discovery, enzymology and engineering. Natural Product Reports 2021, 38, 130–239. DOI: 10.1039/D0NP00027B.

(17) Arnison, P.; Bibb, M.; Bierbaum, G.; Bowers, A.; Bugni, T.; Bulaj, G.; Camarero, J.; Campopiano, D.; Challis, G.; Clardy, J.;, et al. Ribosomally synthesized and post-translationally modified peptide natural products: overview and recommendations for a universal nomenclature. Natural Product Reports 2013, 30, 108–160. DOI: 10.1039/C2NP20085F.

(18) Ayikpoe, R. S.; Zhu, L.; Chen, J. Y.; Ting, C. P.; van der Donk, W. A. Macrocyclization and backbone rearrangement during RiPP biosynthesis by a SAM-dependent domain-of-unknown-function 692. ACS Central Science 2023, 9 (5), 1008–1018. DOI: 10.1021/acscentsci.3c00160.

(19) Manley, O. M.; Shriver, T. J.; Xu, T.; Melendrez, I. A.; Palacios, P.; Robson, S. A.; Guo, Y.; Kelleher, N. L.; Ziarek, J. J.; Rosenzweig, A. C. A multi-iron enzyme installs copper-binding oxazolone/thioamide pairs on a nontypeable *Haemophilus influenzae* virulence factor. Proceedings of the National Academy of Sciences 2024, 121 (28), e2408092121. DOI: 10.1073/pnas.2408092121.

(20) Leprevost, L.; Jünger, S.; Lippens, G.; Guillaume, C.; Sicoli, G.; Oliveira, L.; Rivera-Millot, A.; Billon, G.; Henry, C.; Antoine, R.;, et al. A widespread family of ribosomal peptide metallophores involved in bacterial adaptation to metal stress. Proceedings of the National Academy of Sciences 2024, 121 (49), e2408304121. DOI: 10.1073/pnas.2408304121.

(21) Tang, Y.; Zhong, W.; Fu, L.; Asante, E.; Kostenko, A.; Vidya, F.; Mandelare-Ruiz, P.; Adeogun, T. T.; Anderson, G. P.; Edmonds, B. E.;, et al. Phylogenomic identification of a highly conserved copper-binding RiPP biosynthetic gene cluster in marine *Microbulbifer Bacteria*. ACS Chemical Biology 2025, 20 (10), 2462–2474. DOI: 10.1021/acschembio.5c00507.

(22) Ting, C. P.; Funk, M. A.; Halaby, S. L.; Zhang, Z.; Gonen, T.; Van Der Donk, W. A. Use of a scaffold peptide in the biosynthesis of amino acid–derived natural products. Science 2019, 365 (6450), 280–284. DOI: 10.1126/science.aau6232.

(23) Yu, Y.; van der Donk, W. A. Biosynthesis of 3-thia-á-amino acids on a carrier peptide. Proceedings of the National Academy of Sciences 2022, 119 (29), e2205285119. DOI: 10.1073/pnas.2205285119.

(24) Padhi, C.; Zhu, L.; Chen, J. Y.; Huang, C.; Moreira, R.; Challis, G. L.; Cryle, M. J.; van der Donk, W. A. Biosynthesis of biphenomycin-like macrocyclic peptides by formation and cross-linking of *ortho*-tyrosines. Journal of the American Chemical Society 2025, 147 (27), 23781–23796. DOI: 10.1021/jacs.5c06044.

(25) Strunk, E.; Lobert, A.; Khorovich, T.; Lucio, K. G.; Richarz, R.; Hohmann, M.; D’Agostino, P.; Gulder, T. Biosynthesis of the biphenomycin family of potent antibiotics. Angewandte Chemie International Edition 2025, 64 (51), e202516156. DOI: 10.1002/anie.202516156.

(26) Ouyang, Y.; Yu, Y.; Zhu, L.; Nguyen, D. T.; van der Donk, W. A. Oxidative peptide backbone cleavage by a HEXXH enzyme during RiPP biosynthesis. Journal of the American Chemical Society 2026, 148 (3), 3551–3561. DOI: 10.1021/jacs.5c19246.

(27) Hebron, D. P.; Shriver, T. J.; Ziarek, J. J.; Rosenzweig, A. C. Bis-hydroxylation of Homocitrulline Catalyzed by a Multinuclear Nonheme Iron-Dependent Oxidative Enzyme during RiPP Biosynthesis. *bioRxiv* 2026. DOI: 10.64898/2026.05.01.722236.

(28) Chioti, V. T.; Clark, K. A.; Ganley, J. G.; Han, E. J.; Seyedsayamdost, M. R. N–Cá bond cleavage catalyzed by a multinuclear iron oxygenase from a divergent methanobactin-like RiPP gene cluster. Journal of the American Chemical Society 2024, 146 (11), 7313–7323. DOI: 10.1021/jacs.3c11740.

(29) Nguyen, D. T.; Zhu, L.; Mitchell, D. A.; van der Donk, W. A. Biosynthesis of macrocyclic peptides with C-terminal β-amino-α-keto acid groups by three different metalloenzymes. ACS Central Science 2024, 10 (5), 1022–1032. DOI: 10.1021/acscentsci.4c00088.

(30) Ahearn, C.; Kirkham, C.; Chaves, L.; Kong, Y.; Pettigrew, M.; Murphy, T. Discovery and contribution of nontypeable *Haemophilus influenzae* NTHI1441 to human respiratory epithelial cell invasion. Infection and Immunity 2019, 87 (11). DOI: 10.1128/iai.00462-19.

(31) Burkhart, B. J.; Hudson, G. A.; Dunbar, K. L.; Mitchell, D. A. A prevalent peptide-binding domain guides ribosomal natural product biosynthesis. Nature Chemical Biology 2015, 11, 564–570. DOI: 10.1038/nchembio.1856.

(32) Manley, O. M.; Shriver, T. J.; Ayala, J. M.; Owen, B. C.; Ziarek, J. J.; Rosenzweig, A. C. Differentiating 5-thiooxazoles from oxazolone-coupled thioamides in RiPP natural products. *bioRxiv* 2026. DOI: 10.64898/2026.06.05.730506.

(33) Gadgil, M. G.; Dommaraju, S. R.; Liu, X.; Battiste, A. J.; Bregman, M. H.; Mitchell, D. A. Biosynthesis of peptidic thiooxazole metallophores installed by multinuclear nonheme iron enzymes. ACS Chemical Biology 2025, 21 (4), 722–732. DOI: 10.1021/acschembio.5c00987.

(34) Stookey, L. L. Ferrozine – a new spectrophotometric reagent for iron. Analytical Chemistry 1970, 42 (7), 779–781. DOI: 10.1021/ac60289a016.

(35) Davoust, C. E.; Doan, P. E.; Hoffman, B. M. Q-band pulsed electron spin-echo spectrometer and its application to ENDOR and ESEEM. Journal of Magnetic Resonance, Series A 1996, 119 (1), 38–44. DOI: 10.1006/jmra.1996.0049.

(36) Epel, B.; Gromov, I.; Stoll, S.; Schweiger, A.; Goldfarb, D. Spectrometer manager: A versatile control software for pulse EPR spectrometers. Concepts in Magnetic Resonance Part B: Magnetic Resonance Engineering: An Educational Journal 2005, 26 (1), 36–45. DOI: 10.1002/cmr.b.20037.

(37) Petasis, D. T.; Hendrich, M. P. Quantitative interpretation of multifrequency multimode EPR spectra of metal containing proteins, enzymes, and biomimetic complexes. In Methods in Enzymology, Vol. 563; Elsevier, 2015; pp 171–208. DOI: 10.1016/bs.mie.2015.06.025.

(38) Jünger, S.; Leprevost, L.; Zirah, S.; Carassus, C.; Arnoux, P.; Antoine, R.; Jacob-Dubuisson, F.; Li, Y. Sec Signal Peptide Doubles Up as a Leader Sequence in Bufferin Biosynthesis. ACS Chemical Biology 2026, 21 (6), 1309–1318. DOI: 10.1021/acschembio.6c00045.

(39) Zheng, Y.; Xu, X.; Fu, X.; Zhou, X.; Dou, C.; Yu, Y.; Yan, W.; Yang, J.; Xiao, M.; van der Donk, W. A.;, et al. Structures of the holoenzyme TglHI required for 3-thiaglutamate biosynthesis. Structure 2023, 31 (10), 1220–1232. DOI: 10.1016/j.str.2023.08.004.

(40) Davydov, R. M.; Ménage, S.; Fontecave, M.; Gräslund, A.; Ehrenberg, A. Mixed-valent *μ*-oxo-bridged diiron complexes produced by radiolytic reduction at 77 K studied by EPR. Journal of Biological Inorganic Chemistry 1997, 2, 242–255. DOI: 10.1007/s007750050130.

(41) Noodleman, L.; Peng, C.; Case, D.; Mouesca, J.-M. Orbital interactions, electron delocalization and spin coupling in iron-sulfur clusters. Coordination Chemistry Reviews 1995, 144, 199–244. DOI: 10.1016/0010-8545(95)07011-L.

(42) Joint Center for Structural Genomics (JCSG). Crystal structure of a duf692 family protein (hs_1138) from Haemophilus somnus 129pt at 2.20 A resolution. 2008. DOI: 10.2210/pdb3bww/pdb.

(43) Antoine, R.; Leprevost, L.; Jünger, S.; Zirah, S.; Lippens, G.; Li, Y.; Dubiley, S.; Jacob-Dubuisson, F. Multinuclear non-heme iron dependent oxidative enzymes: Landscape of their substrates, partner proteins and biosynthetic gene clusters. Microbial Genomics 2025, 11 (7), 001462. DOI: 10.1099/mgen.0.001462.

