## Supporting Information for "Thiooxazole Formation on a Nontypeable *Haemophilus influenzae* Virulence Factor Requires a Mixed-Valent Diiron Cofactor"

**Table S1.** Nucleotide sequence of the fusion of HvfBC used in this study. Restriction sites used to clone into pET28a(+)-TEV are underlined.

| HvfBC$_{fusion}$ |
|---|
| CATATGAAATTACAAGGTGCTGGGCTAGGATATCGCCGTAATCTGGCGGAAGATTTCCTGCAG |
| CTGCCGAGCAATAACGCAATCCAGTTCATCGAAGTTGCTCCGGAAAACTGGTCCAAAATGGGT |
| GGCATGGCACGTTATCAATTTGATCAAGCAGCTGAGCGCTTTCCGTTAGCCGTTCATGGCCTGT |
| CTCTGTCCCTGGGCGGTCAGGCACCGCTGGACAGAGAATTGCTGCGTAACACCAAAGCGCTGA |
| TCAACCAGTATAACTCTAGCTTTTTCAGCGAACACTTGTCGTACTGCGAGTGCGAGGGTCACCT |
| GTACGACCTGCTGCCGATGCCGTTCACCGAAGAAGCGGTTAAGCACGTGGCCCAACGTATTCG |
| CGACGTACAGGATTTCCTCGGCCTGCAAATCAGCTTGGAGAACACCTCGTACTACCTGCATTCC |
| CCGACGTCCACCATGAATGAGGTGGAGTTCTTAAATGCCATCGCGCAAGAAGCGGACTGTGGT |
| ATTCATCTGGACGTGAACAACATTTATGTTAATGGTGTTAACCACGGCCTGCTGGACCCGTATA |
| TTTTTTTGGACCAGGTCGATGTGAAGCGCGTGAACTATATCCACATCGCCGGTCACGATGAAG |
| AGCACAGCGCTGCGCAAGTTGTTGAGAACAGCGCTAATGAGAGCTTCAATAAAGTGAAGGGC |
| GCATATCGTCATCTGCCGGAGTTGTTAATTGATACCCATGGTGAAGCGGTGAAGGGTACTGTTT |
| GGGATTTGTTGGAGTATGCGTACCAGCGTCTGCCAACCATTCCGCCTACGTTGCTTGAGCGTGA |
| CTTCAACTTTCCACCGTTTGCAGAGCTCTACGCCGAAGTGGAGCACATCGCTCAACTGCAGCA |
| GAAATACGCGCATACCGAAGTCATGAGCTACGCGGCGCTACCCAAGAGTTCATTAAAAGAAA |
| CACAACAAGCGCTGGCGAATGCGATCAGATTAGGTAATGCGGATCCATTGAATGGCTATGCAG |
| CCAGCCGTCTTGCGGTCTATACCCGTCTGGTTCGCAACAACGCGTTTGGCTTTATTGATCGCTG |
| CTTTGTGGAAGCACCGCTGCATATCGAACCGGAATATTGGAAAAACGCTAAAGAGAACTTCGT |
| CCAAAACGGCAACGCTCACTCTCCGTATTTCCAGGATATCGCCGGTGAATTTCTCTTGTTTTGT |
| CAGGAGAAAGAGATTTTCGACACCAATATTCTGGCGCTGATGGATTTCGAGAACACCCAGCTG |
| CTGGCTGAGGTTAGCTTGGCTAAGGTGCCGGAGAAGTTCGAGTGGAATCGTCACTCCGTGATG |
| CAGCTGAGCGGTGCAGCATACCTCAAGTCCTACGACGTTGATTTCTTGTCTAGCGACTTCAAGC |
| AGTTTGACGACACTCCGATCCAGGCGATTATCTGGCGTGATTCGGACTTCCGCATCCAACAGC |
| AAATTCTGTCCGAACTGGACTACTGGCTGTTAAGCTACCTGCAAGAACAGCCGAACAGCCTGG |
| AAAACGTGCTGAGCGCGCTGAATACGATGGTGGAAGATAGCACCTCTATTGTTCCGTTGCTGG |
| AACAGGTTTGGATGAAATGGGTTACCAGCGAGGTGATCTACCCGGAGCAACGTTAAAAGCTT |

**Table S2.** Primers used for site-directed mutagenesis of HvfBC$_{fusion}$. The mutated codon is bolded, and the changed nucleotides are underlined. The nonoverlapping region is italicized.

| Mutation | Direction | Sequence (5'→3') |
|---|---|---|
| N175A | forward | GGACGTGAAC**GCC**ATTTATG*TTAATGGTGTTAACCACGGCC* |
| | reverse | CATAAAT**GGC**GTTCACGTCC*AGATGAATACCACAGTCCGC* |
| N175S | forward | GGACGTGAAC**AGC**ATTTATG*TTAATGGTGTTAACCACGGCC* |
| | reverse | CATAAAT**GCT**GTTCACGTCC*AGATGAATACCACAGTCCGC* |
| D242A | forward | GTTAATT**GCT**ACCCATGGTG*AAGCGGTGAAGGGTACTGTTTGG* |
| | reverse | CACCATGGGT**AGC**AATTAAC*AACTCCGGCAGATGACGATATGC* |
| D242S | forward | GTTAATT**TCT**ACCCATGGTG*AAGCGGTGAAGGGTACTGTTTGG* |
| | reverse | CACCATGGGT**AGA**AATTAAC*AACTCCGGCAGATGACGATATGC* |
| H244A | forward | GTTAATTGATACC**GCT**GGTG*AAGCGGTGAAGGGTACTGTTTGG* |
| | reverse | CACC**AGC**GGTATCAATTAAC*AACTCCGGCAGATGACGATATGC* |
| H244S | forward | GTTAATTGATACC**TCT**GGTG*AAGCGGTGAAGGGTACTGTTTGG* |
| | reverse | CACC**AGA**GGTATCAATTAAC*AACTCCGGCAGATGACGATATGC* |

**Table S3.** Masses of HvfA peptides following reactions with $HvfBC_{coexp}$ or reactions omitting individual reaction components. The mass difference (Δm) represents the difference between the detected mass and the theoretical unmodified mass (10281.7 Da for the full-length HvfA and 7981.9 Da for HvfA lacking the signal peptide). The number of modifications represents the approximate average number of post-translational modifications (–4 Da each) per peptide.

| | Full-length peptide | | | Signal peptide cleaved | | |
|---|---|---|---|---|---|---|
| | Detected mass (Da) | Δm (Da) | # of mod. | Detected mass (Da) | Δm (Da) | # of mod. |
| $HvfBC_{coexp}$ | 10260.6 ± 1.9 | –21.1 | 5.3 | 7978.7 ± 2.9 | –3.2 | 0.8 |
| –HvfBC | 10281.1 ± 0.3 | –0.6 | 0.2 | 7981.4 ± 0.4 | –0.5 | 0.1 |
| –DTT | 10279.2 ± 2.0 | –2.5 | 0.6 | 7979.2 ± 2.1 | –2.7 | 0.7 |
| $–O_2$ | 10280.3 ± 0.5 | –1.4 | 0.3 | 7981.7 ± 0.4 | –0.2 | 0.0 |

**Table S4.** Masses of HvfA peptides following reactions with $HvfBC_{coexp}$, HvfC (omitting HvfB), HvfB (omitting HvfC), $HvfBC_{comb}$, and $HvfBC_{fusion}$. The mass difference (Δm) represents the difference between the detected mass and the theoretical unmodified mass (10281.7 Da for full-length HvfA and 7981.9 Da for HvfA lacking the signal peptide). The number of modifications represents the approximate average number of post-translational modifications (–4 Da each) per peptide.

| | Full-length peptide | | | Signal peptide cleaved | | |
|---|---|---|---|---|---|---|
| | Detected mass (Da) | Δm (Da) | # of mod. | Detected mass (Da) | Δm (Da) | # of mod. |
| $HvfBC_{coexp}$ | 10260.6 ± 1.9 | –21.1 | 5.3 | 7978.7 ± 2.9 | –3.2 | 0.8 |
| –HvfB | 10280.8 ± 0.2 | –0.9 | 0.2 | 7981.0 ± 0.2 | –0.9 | 0.2 |
| –HvfC | 10281.1 ± 0.4 | –0.6 | 0.1 | 7981.2 ± 0.4 | –0.7 | 0.2 |
| $HvfBC_{comb}$ | 10280.7 ± 0.2 | –1.0 | 0.2 | 7981.2 ± 0.2 | –0.7 | 0.2 |
| $HvfBC_{fusion}$ | 10257.4 ± 0.6 | –24.3 | 6.0 | 7967.8 ± 0.2 | –14.1 | 3.5 |

**Table S5.** Masses of HvfA peptides following reactions with $HvfBC_{fusion}$ or reactions omitting individual reaction components. The mass difference (Δm) represents the difference between the detected mass and the theoretical unmodified mass (10281.7 Da for the full-length HvfA and 7981.9 Da for HvfA lacking the signal peptide). The number of modifications represents the approximate average number of post-translational modifications (–4 Da each) per peptide.

| | Full-length peptide | | | Signal peptide cleaved | | |
|---|---|---|---|---|---|---|
| | Detected mass (Da) | Δm (Da) | # of mod. | Detected mass (Da) | Δm (Da) | # of mod. |
| $HvfBC_{fusion}$ | 10257.4 ± 0.6 | –24.3 | 6.0 | 7967.8 ± 0.2 | –14.1 | 3.5 |
| –DTT | 10281.1 ± 0.2 | –0.6 | 0.1 | 7981.7 ± 0.2 | –0.2 | 0.1 |
| $–O_2$ | 10280.1 ± 0.7 | –1.6 | 0.4 | 7981.8 ± 0.2 | –0.1 | 0.0 |

**Table S6.** Parameters used to simulate the 4.2-K Mössbauer spectra shown in Figure 5B.

| Spin | Site | δ (mm/s) | $\Delta E_Q$ (mm/s) | η [2] | $A_x/g_n\beta_n$, $A_y/g_n\beta_n$, $A_z/g_n\beta_n$ (T) | % Iron | Assignment |
|---|---|---|---|---|---|---|---|
| 5/2 [1] | 1 | 0.55 | -0.57 | 3 | +15.9, +15.9, +15.9 | 18 | Triferric |
| | 2 | 0.50 | 0.59 | 0 | -17.6, -19.4, -19.4 | 18 | |
| | 3 | 0.46 | -0.40 | -3 | -18.2, -18.2, -19.5 | 18 | |
| 0 | 4 | 0.55 | 0.51 | 1 | 0, 0, 0 | 14 | Diferric |
| | 5 | 0.53 | 1.14 | 0 | 0, 0, 0 | 14 | |
| 1/2 [3] | 6 | 0.49 | -1.11 | 7.2 | -43.1, -55.8, -53.4 | 10 | Fe(II)/Fe(III) |
| | 7 | 1.12 | 2.68 | 0.3 | 26.5, 22.5, 18.0 | 10 | |

[1] The current Mössbauer simulations use an $S = 5/2$ spin Hamiltonian to simulate the triferric cofactor with the individual iron sites listed in the table as Sites 1-3. The spin Hamiltonian parameters used for the current simulation are $D = 0.3$ cm$^{-1}$, $E/D = 0.2$, g = [2, 2, 2].
[2] The asymmetric parameter of quadrupole splitting, η, normally is defined with a value range between 0 and 1 if the three principal components of the EFG tensor are chosen in the following order. $|V_{xx}|\leq|V_{yy}|\leq|V_{zz}|$. Here η > 1 or < -1 indicates that the largest component of the EFG tensor is oriented away from the *z* axis defined by the zero field splitting tensor.
[3] The simulation parameters for the Fe(II)/Fe(III) mixed-valent cofactor are from ref[1], the A values are slightly adjusted to cover the extend of magnetic splittings of this species observed in the spectrum recorded under a 7 T magnetic field parallel applied to the gamma radiation.

**Table S7.** Masses of HvfA peptides following reactions with $HvfBC_{fusion}$ variants with ligands to the third iron-binding site mutated. The mass difference (Δm) represents the difference between the detected mass and the theoretical unmodified mass (10281.7 Da for the full-length HvfA and 7981.9 Da for HvfA lacking the signal peptide). The number of modifications represents the approximate average number of post-translational modifications (–4 Da each) per peptide.

| | Full-length peptide | | | Signal peptide cleaved | | |
|---|---|---|---|---|---|---|
| | Detected mass (Da) | Δm (Da) | # of mod. | Detected mass (Da) | Δm (Da) | # of mod. |
| WT | 10257.8 ± 0.3 | –23.9 | 6.0 | 7967.8 ± 0.3 | –14.1 | 3.5 |
| N175A | 10281.6 ± 0.2 | –0.1 | 0.0 | 7981.7 ± 0.2 | –0.2 | 0.0 |
| D242A | 10281.6 ± 0.1 | –0.1 | 0.0 | 7981.7 ± 0.2 | –0.2 | 0.0 |
| H244A | 10281.3 ± 0.2 | –0.4 | 0.1 | 7981.7 ± 0.1 | –0.2 | 0.0 |
| N175S | 10281.1 ± 0.7 | –0.6 | 0.1 | 7981.5 ± 0.5 | –0.4 | 0.1 |
| D242S | 10281.6 ± 0.2 | –0.1 | 0.0 | 7981.8 ± 0.1 | –0.1 | 0.0 |
| H244S | 10281.4 ± 0.2 | –0.1 | 0.1 | 7981.8 ± 0.1 | –0.1 | 0.0 |

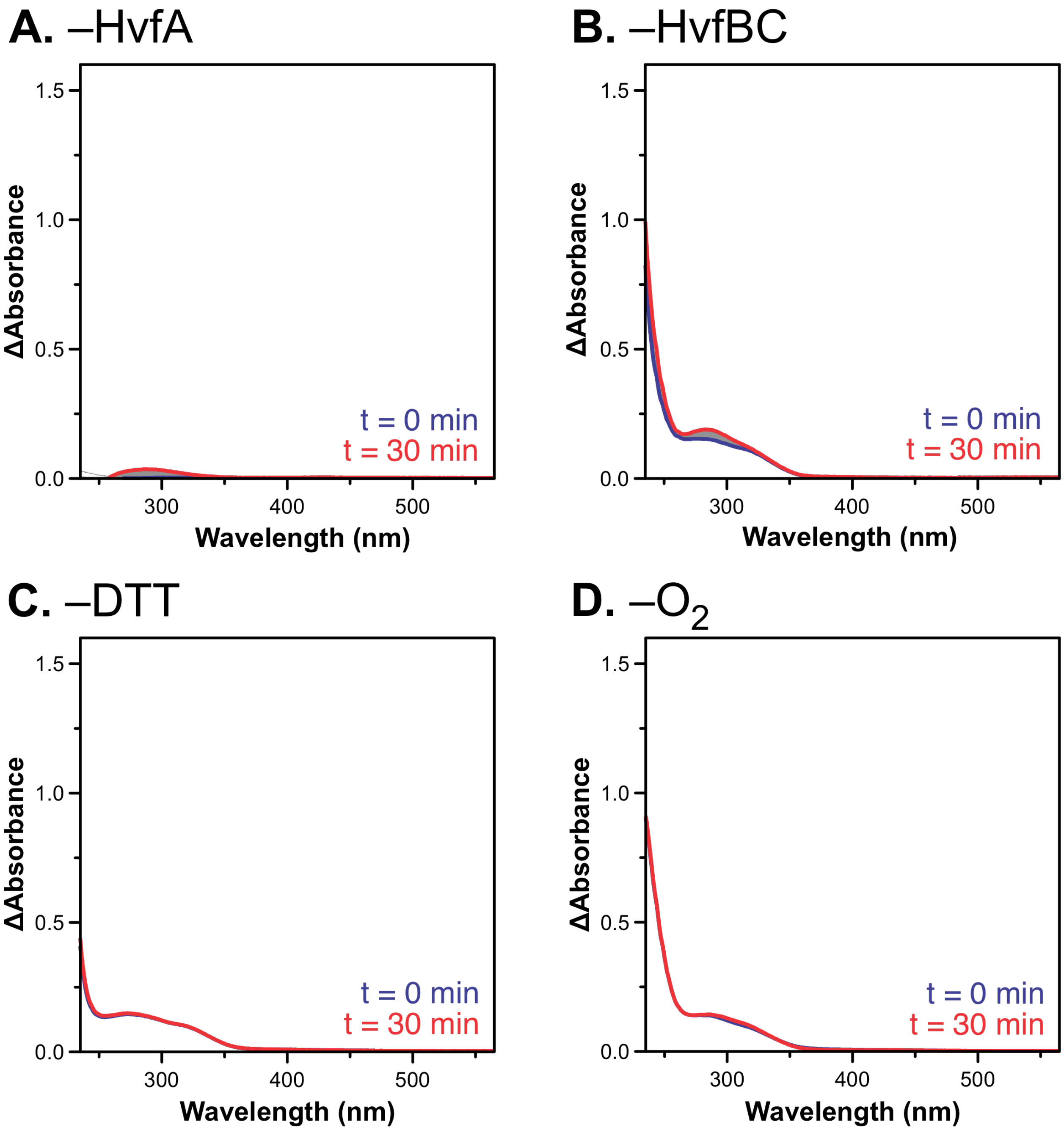

**Figure S1.** Representative control reactions of $HvfBC_{coexp}$ and HvfA omitting HvfA (**A**), $HvfBC_{coexp}$ (**B**), DTT (**C**), and $O_2$ (**D**). UV-visible spectra of the reaction progress, measured every 30 s (gray). The initial spectrum is shown in blue (t = 0 min), and the final absorbance of the reaction is shown in red (t = 30 min). The spectrum of $HvfBC_{coexp}$ was subtracted from the data for clarity. Reactions were performed using 50 µM HvfA, 5 µM $HvfBC_{coexp}$, 1 mM DTT, and 1 mM $O_2$ where applicable.

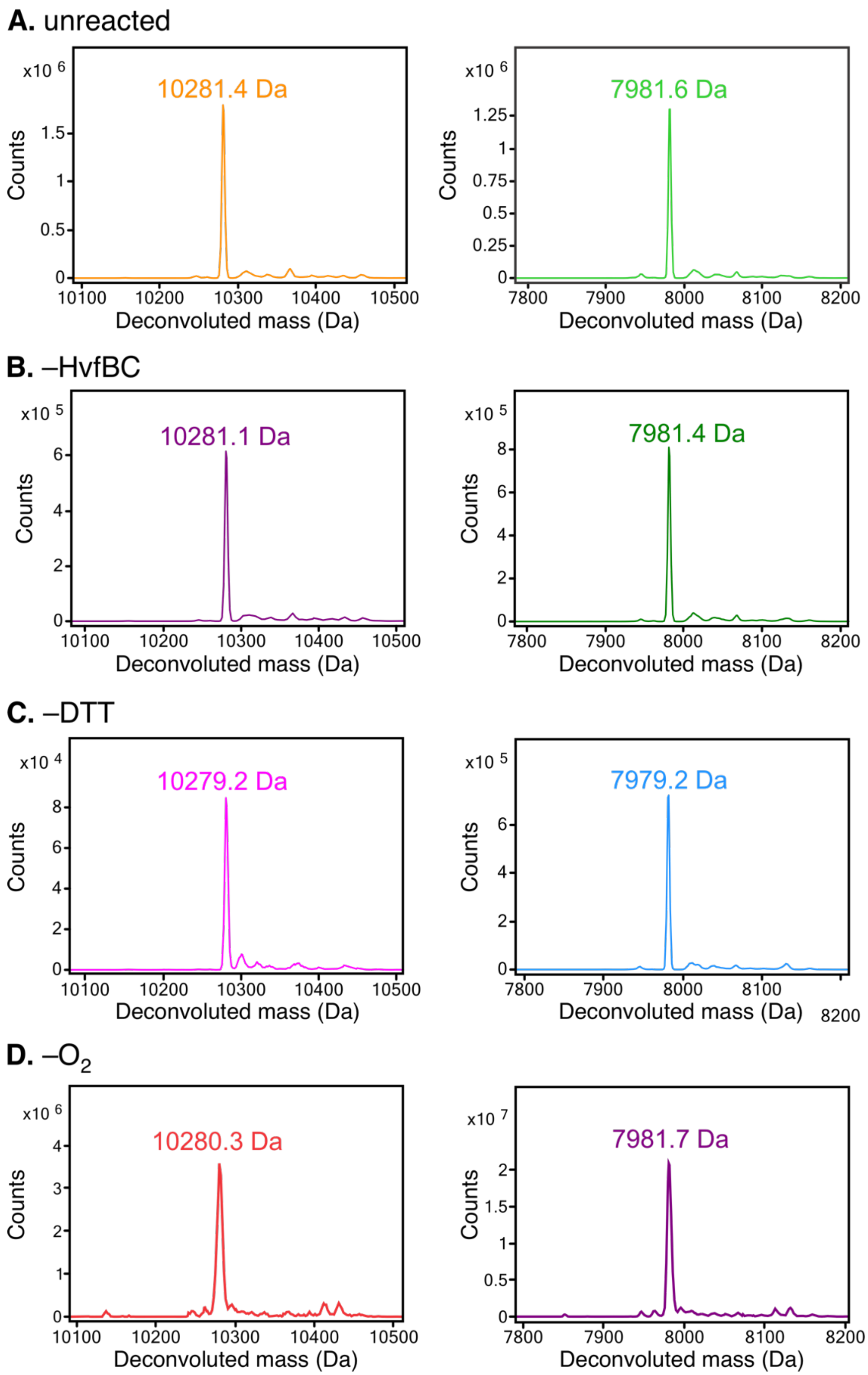

**Figure S2.** Intact protein mass spectrometry of HvfA unreacted (**A**) or following reaction with $HvfBC_{coexp}$ omitting enzyme (**B**), reductant (**C**), or $O_2$ (**D**). Results are shown as deconvoluted mass spectra corresponding to the full-length peptide (left) and that with the signal peptide cleaved (right). Peptides were treated with 1 mM DTT prior to analysis.

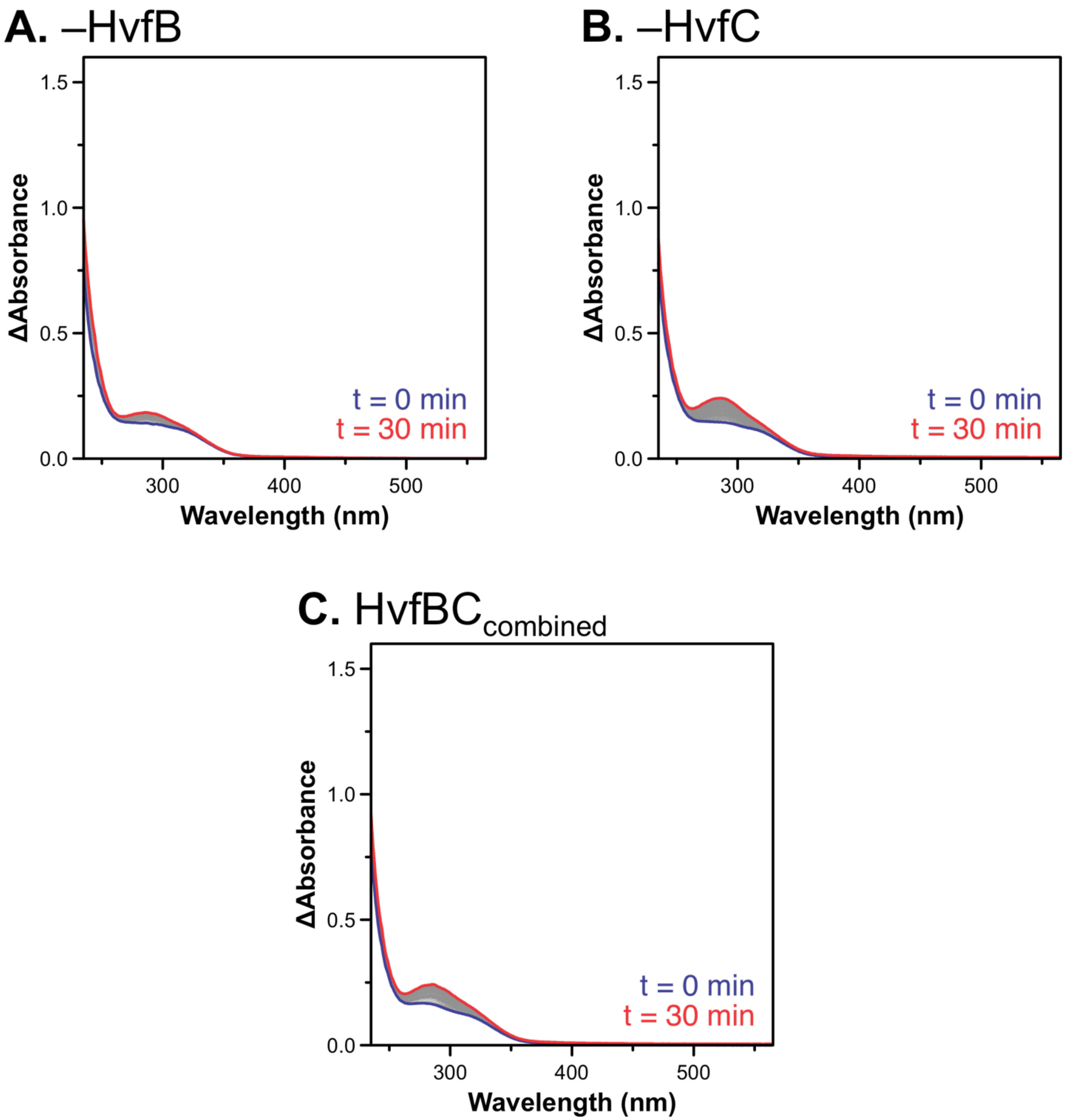

**Figure S3.** Representative reactions of HvfBC and HvfA omitting HvfB (**A**), omitting HvfC (**B**), and combining HvfB and HvfC that were independently purified (**C**). UV-visible spectra of the reaction progress were measured every 30 s (gray). The initial spectrum is shown in blue (t = 0 min), and the final absorbance of the reaction is shown in red (t = 30 min). The spectrum of HvfB and/or HvfC was subtracted from the data for clarity. Reactions were performed using 50 µM HvfA, 5 µM HvfB and/or HvfC, 1 mM DTT, and 1 mM $O_2$.

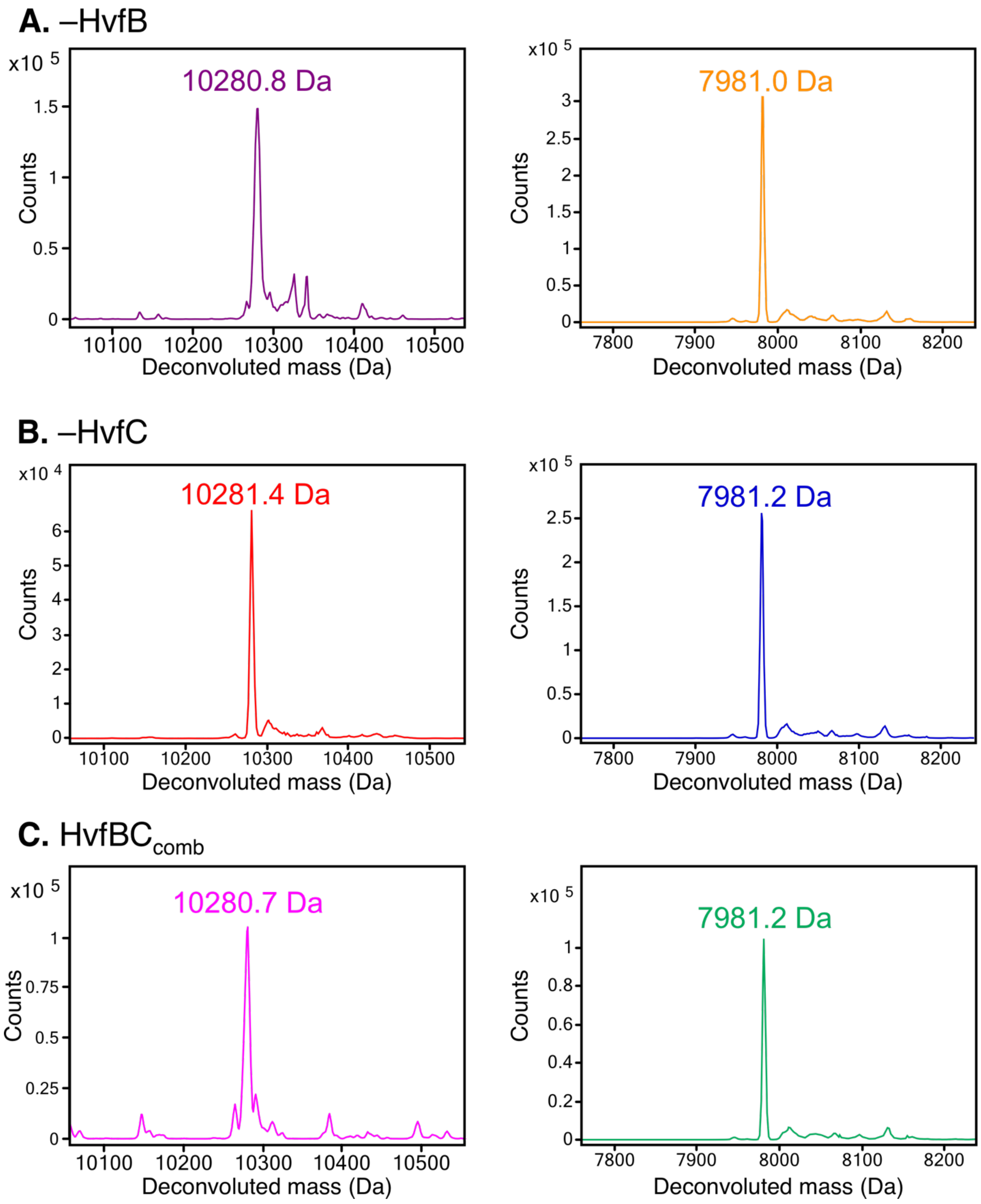

**Figure S4.** Intact protein mass spectrometry of HvfA following reaction with HvfC (omitting HvfB, **A**) HvfB (omitting HvfC, **B**), and HvfBC$_{comb}$ (**C**). Results are shown as deconvoluted mass spectra corresponding to the full-length peptide (left) and that with the signal peptide cleaved (right). Peptides were treated with 1 mM DTT prior to analysis.

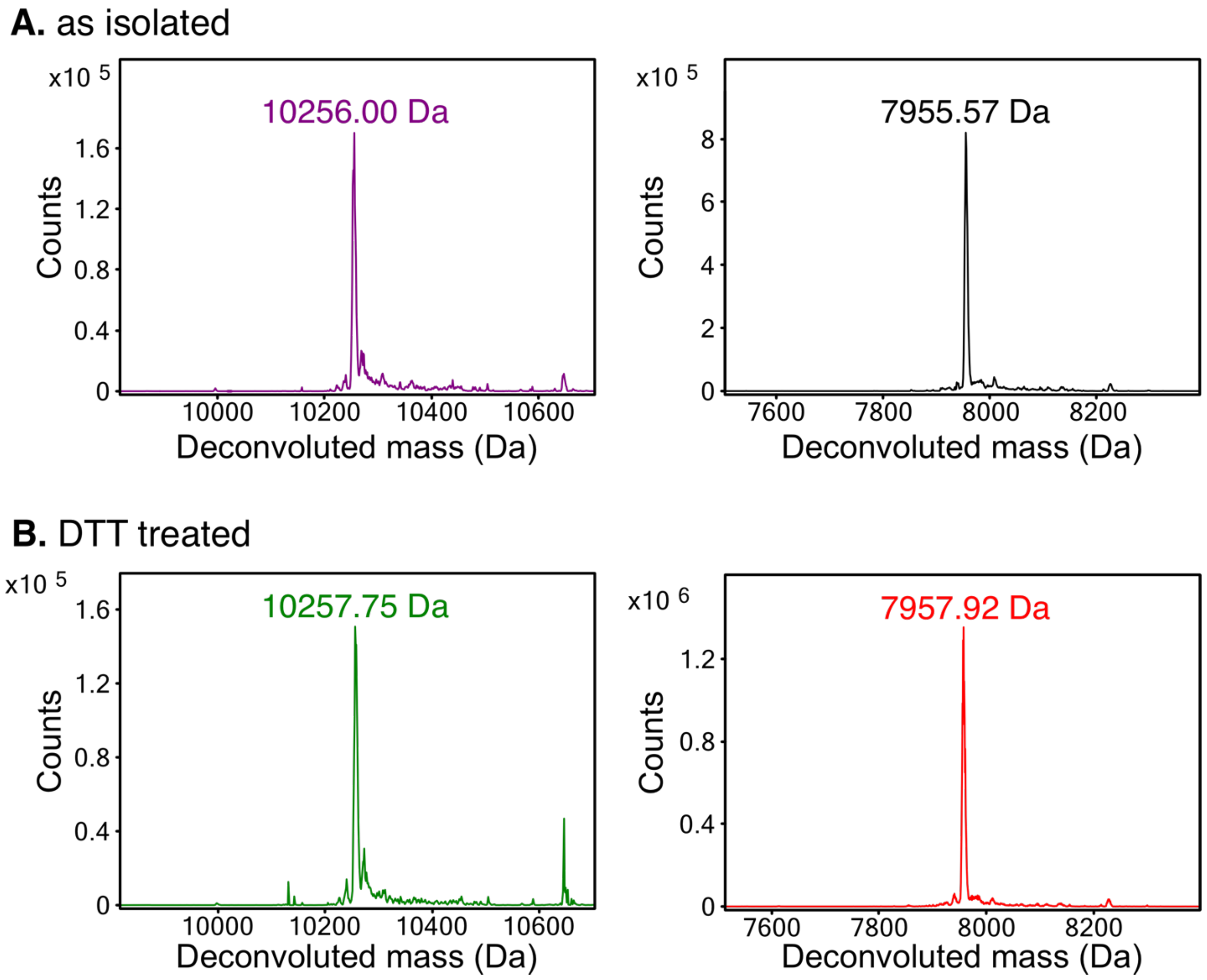

**Figure S5.** Deconvoluted intact protein mass spectra of HvfA after coexpression with $HvfBC_{fusion}$ as isolated (**A**) and upon treatment with DTT (**B**). The full-length peptide is shown on the left and the peptide with the signal-sequence cleaved is shown on the right.

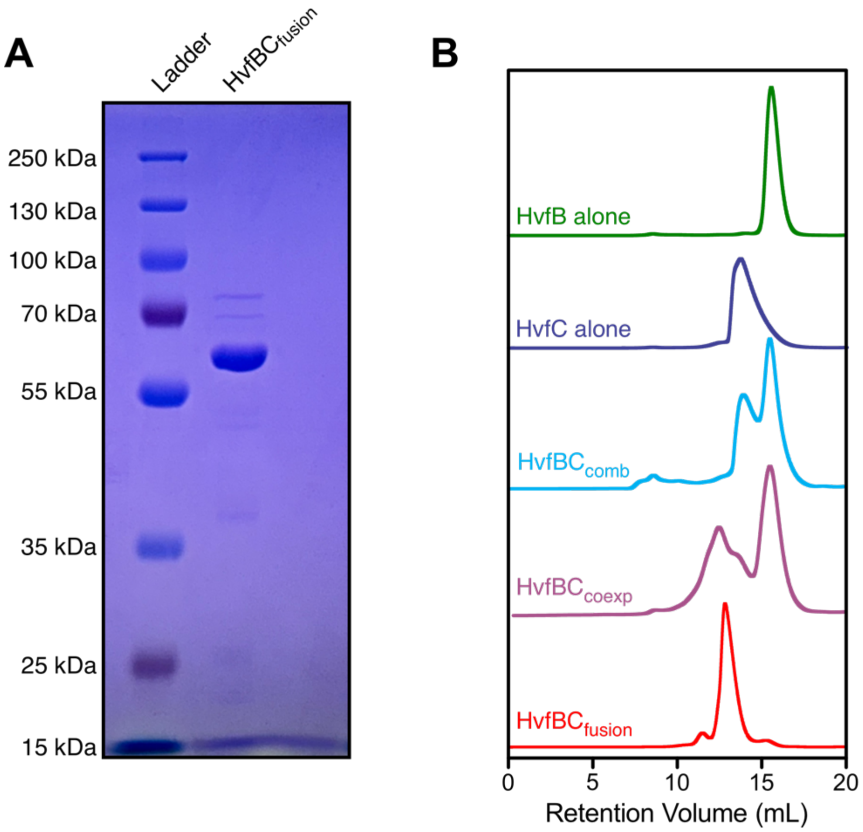

**Figure S6.** (**A**) SDS-PAGE of $HvfBC_{fusion}$ following purification by Ni-affinity chromatography. The calculated molecular weight of $HvfBC_{fusion}$ is 64.6 kDa. (**B**) Size exclusion chromatograms of HvfB, HvfC, $HvfBC_{comb}$, $HvfBC_{coexp}$, and $HvfBC_{fusion}$. The top four chromatograms are taken from ref[2].

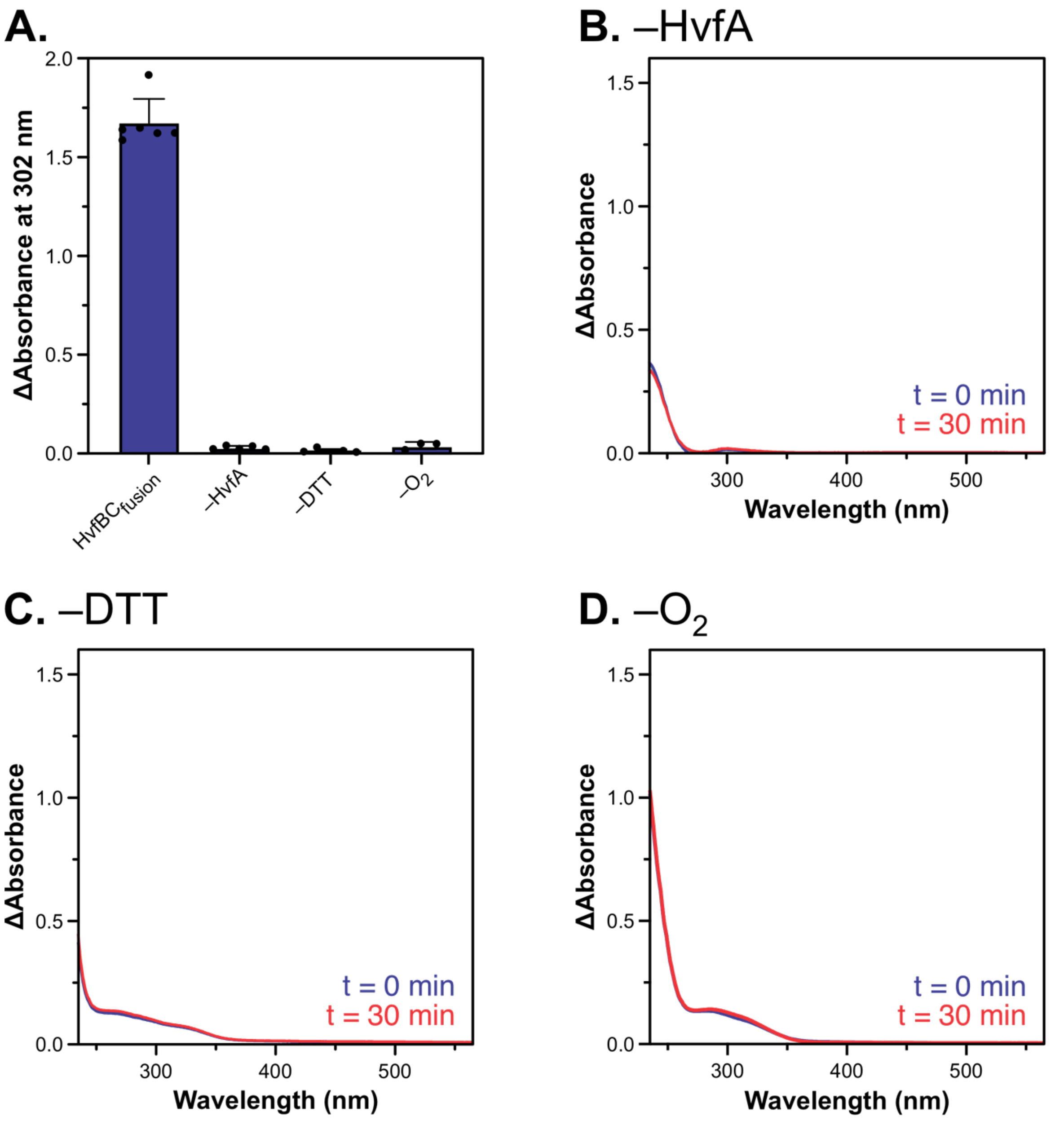

**Figure S7.** Control reactions of $HvfBC_{fusion}$ and HvfA. The increase in absorbance at 302 nm is shown for the reaction of $HvfBC_{fusion}$ and control reactions omitting reaction components (**A**). UV-visible spectra of the reaction progress, measured every 30 s (gray), omitting HvfA (**B**), DTT (**C**), and $O_2$ (**D**). The initial absorbance spectrum is shown in blue (t = 0 min), and the final absorbance spectrum is shown in red (t = 30 min). The spectrum of $HvfBC_{fusion}$ was subtracted from the data for clarity. Reactions were performed using 50 µM HvfA, 5 µM $HvfBC_{fusion}$, 1 mM DTT, and 1 mM $O_2$ where applicable.

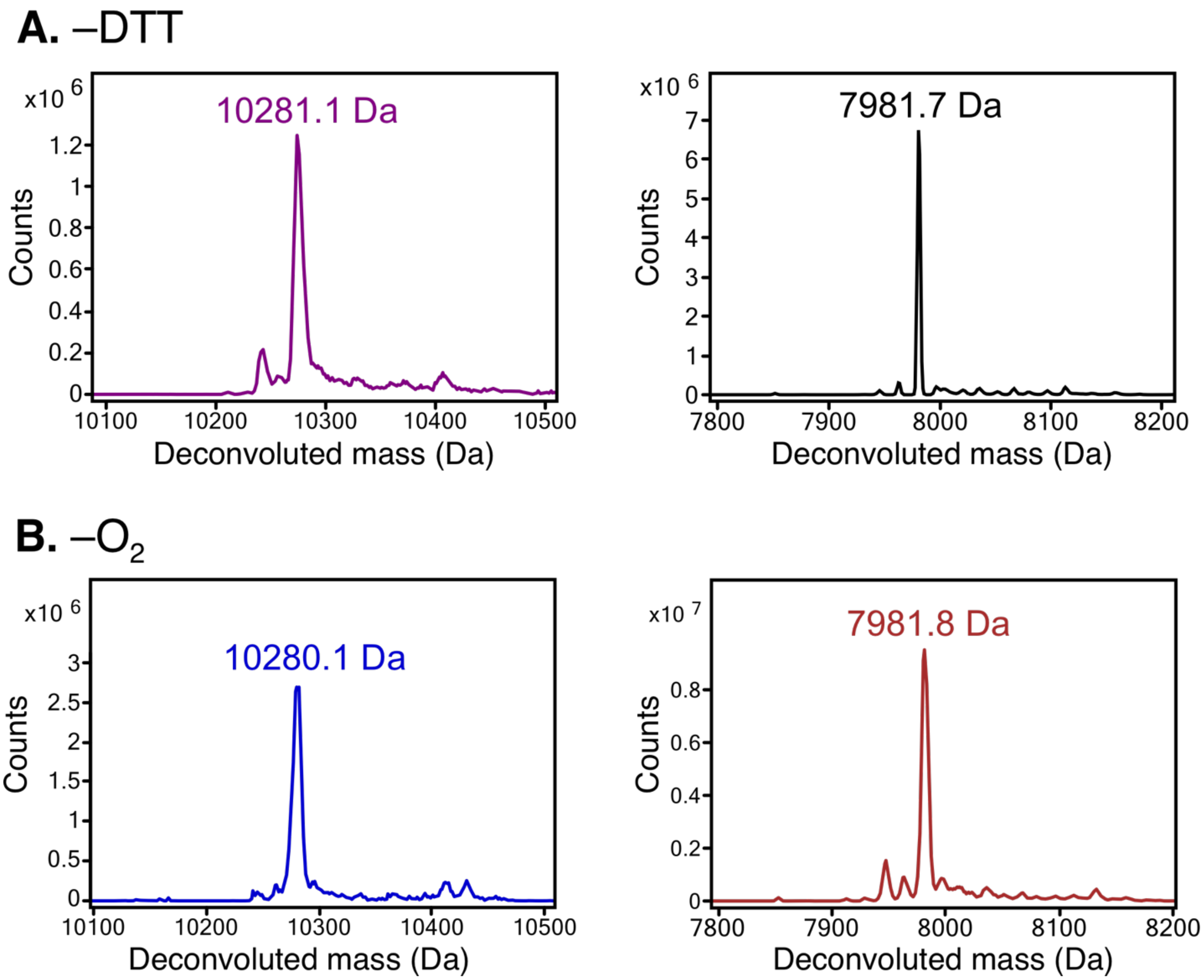

**Figure S8.** Intact protein mass spectrometry of HvfA following reaction with $HvfBC_{fusion}$ omitting reductant (**A**) or oxygen (**B**). Results are shown as deconvoluted mass spectra corresponding to the full-length peptide (left) and that with the signal peptide cleaved (right). Peptides were treated with 1 mM DTT prior to analysis.

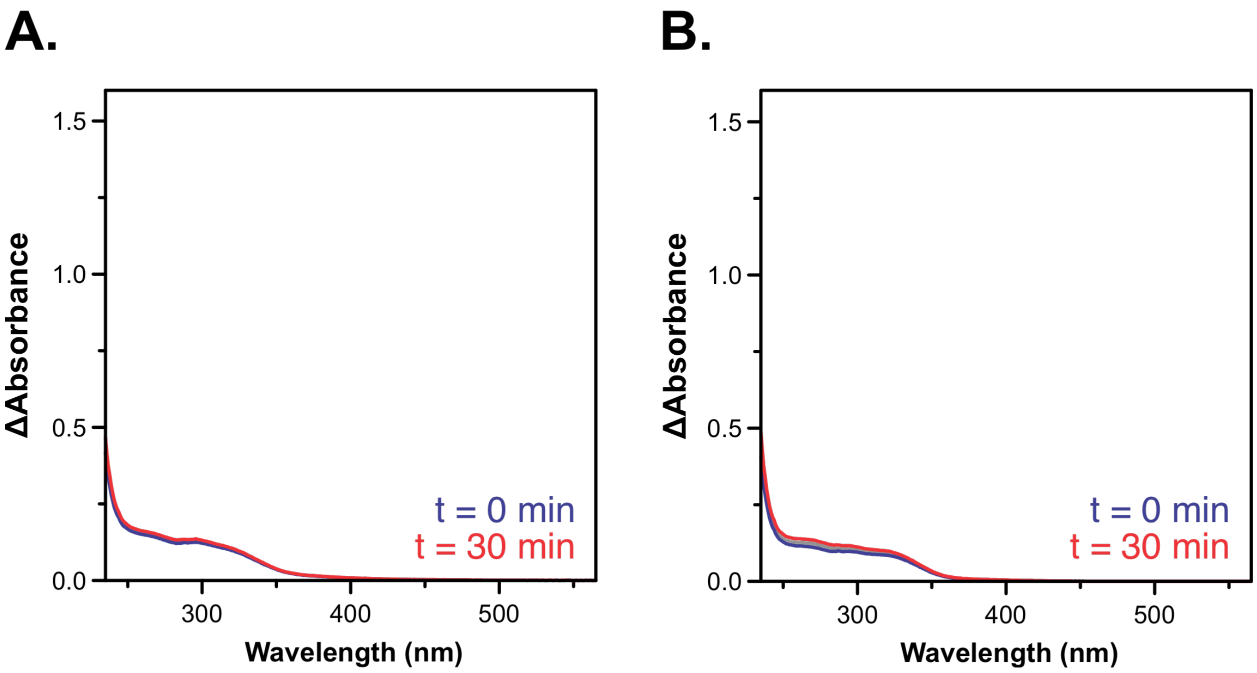

**Figure S9.** The reactions of DTT-treated HvfA with as-isolated $HvfBC_{fusion}$ (**A**) and ascorbate-treated $HvfBC_{fusion}$ with as-isolated HvfA (**B**), without the addition of excess reductant to the reaction. Reactions were monitored by UV-visible spectra measured every 30 s (gray). The initial absorbance spectrum is shown in blue (t = 0 min), and the final absorbance spectrum is shown in red (t = 30 min). The spectrum of $HvfBC_{fusion}$ was subtracted from the data for clarity. Reactions were performed using 50 µM HvfA, 5 µM $HvfBC_{fusion}$, and 1 mM $O_2$.

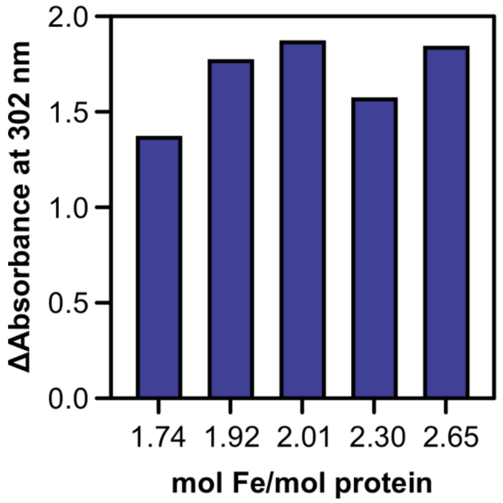

**Figure S10.** Enzymatic activity of $HvfBC_{fusion}$, copurified with different amounts of iron, as measured by the absorbance increase at 302 nm.

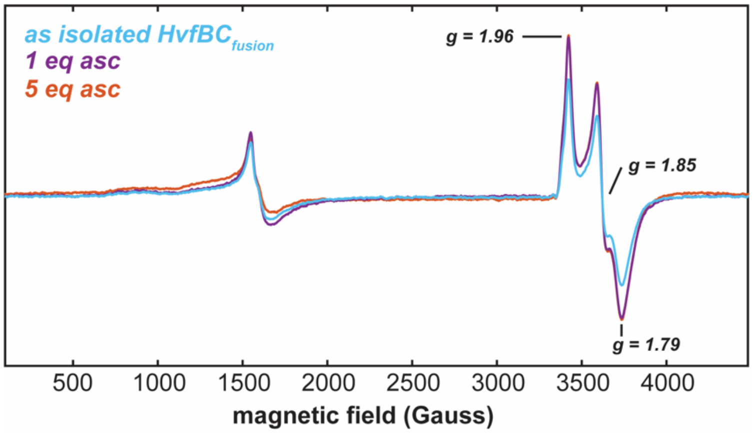

**Figure S11.** X-band EPR spectra of $HvfBC_{fusion}$ with excess sodium ascorbate. Addition of excess ascorbate beyond 1 molar equivalent did not further enhance the $S = 1/2$ signal. *EPR conditions:* microwave frequency ~9.3 GHz, temperature 12 K, 320 ms time constant, 10 G modulation, 25 dB microwave attenuation.

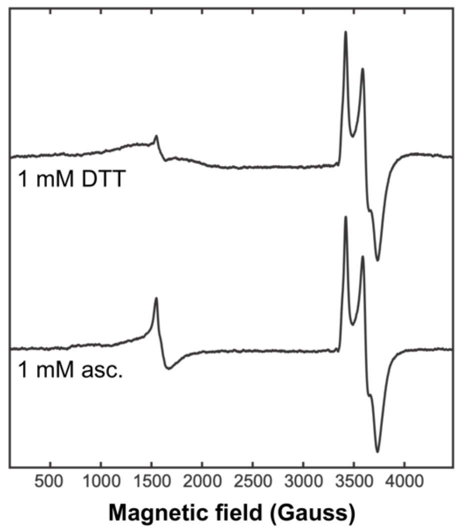

**Figure S12.** X-band EPR spectra of $HvfBC_{fusion}$ treated with 1 mM dithiothreitol (DTT, top) or 1 mM sodium ascorbate (bottom). *EPR conditions:* microwave frequency ~9.3 GHz, temperature 12 K, 320 ms time constant, 10 G modulation, 25 dB microwave attenuation.

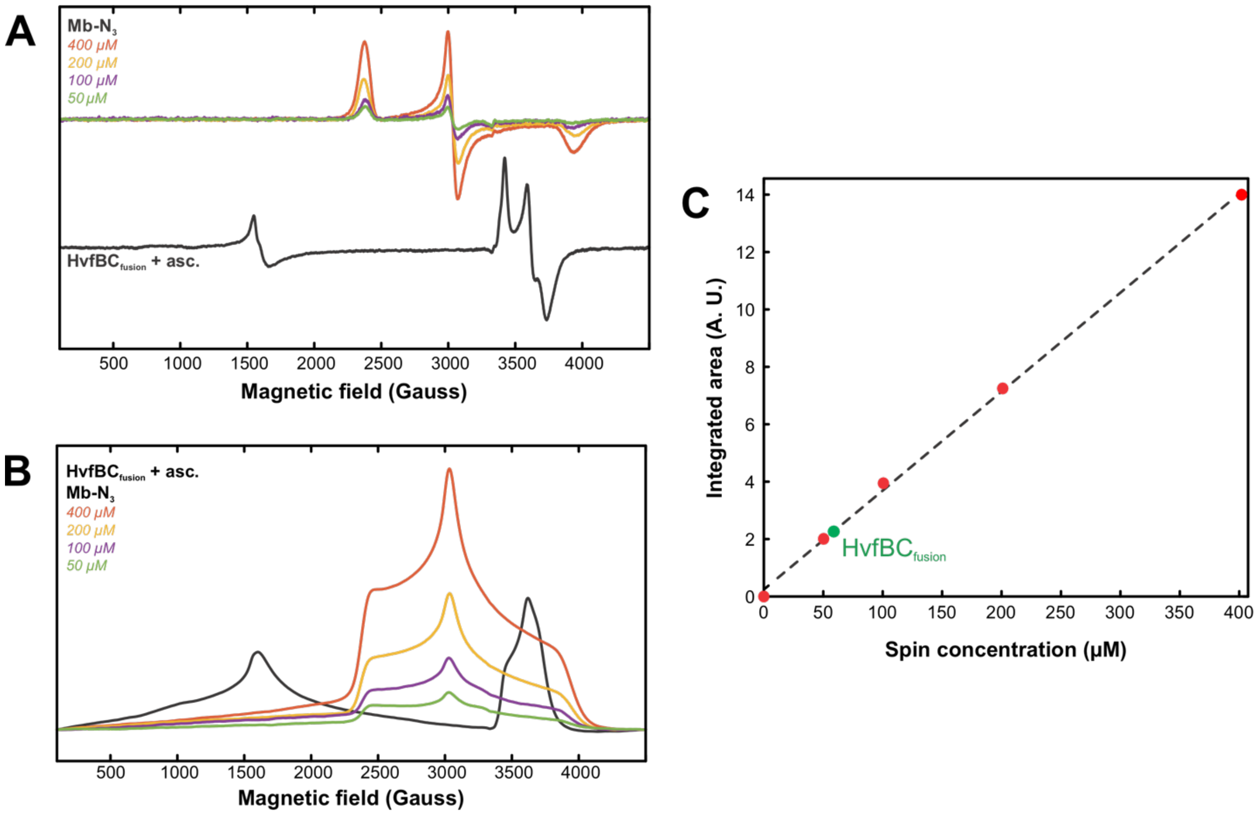

**Figure S13.** Spin quantification of the $S = 1/2$ signal of HvfBC$_{\text{fusion}}$ treated with sodium ascorbate. (**A**) X-band EPR spectra of $N_3$-ligated myoglobin (Mb) at concentrations ranging from 50 to 400 µM and ascorbate-treated HvfBC$_{\text{fusion}}$. (**B**) Integration of the spectra shown in panel **A**. (**C**) A plot of the area under the peaks shown in panel **B** versus concentration. The area of the HvfBC$_{\text{fusion}}$ $g \approx 2$ signal is labeled and shown in green and corresponds to ~60 µM spin per 300 µM protein. *EPR conditions:* microwave frequency ~9.3 GHz, temperature 12 K, 320 ms time constant, 10 G modulation, 25 dB microwave attenuation.

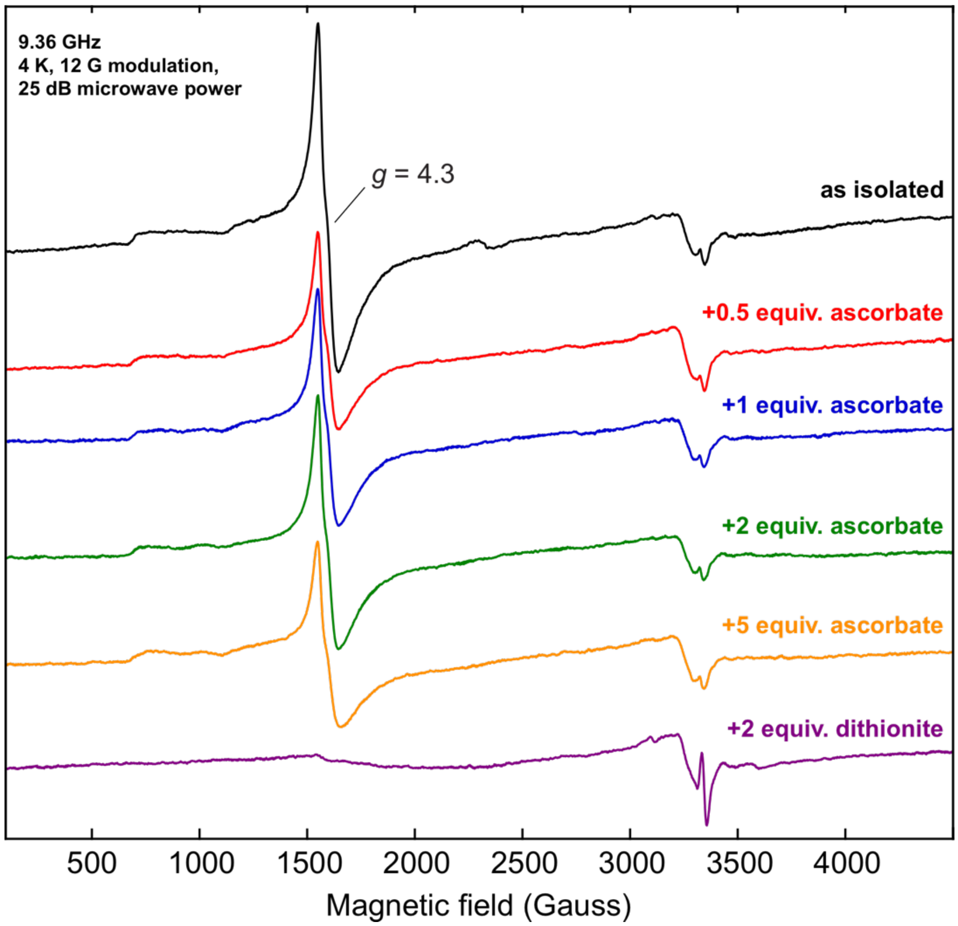

**Figure S14.** X-band EPR spectra of HvfB as isolated, treated with varying molar equivalents of sodium ascorbate, or sodium dithionite under anaerobic atmosphere. The marked signal at $g = 4.3$ is consistent with high-spin $S = 5/2$ Fe (mono/triferric). The small signals at $g = 2.0$ and just above $g = 2.0$ are attributable to mediator dye and contaminating copper, respectively. *EPR conditions:* microwave frequency ~9.3 GHz, temperature 4 K, 320 ms time constant, 12 G modulation, 25 dB microwave attenuation.

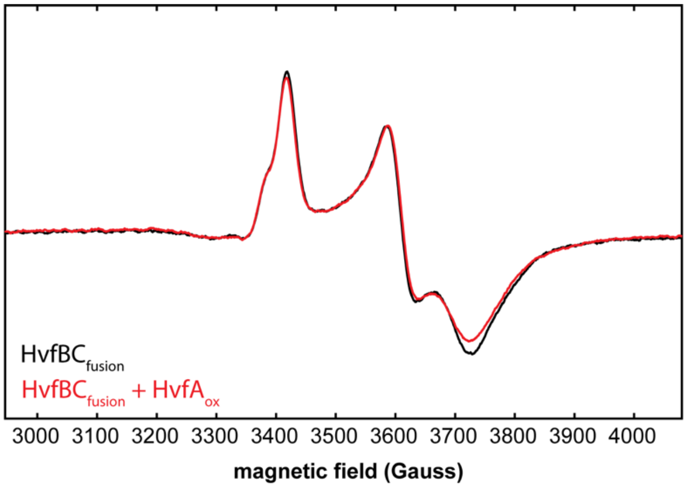

**Figure S15.** Addition of as-isolated HvfA to mixed-valent $HvfBC_{fusion}$ causes no perturbation of the $S$ = 1/2 X-band EPR signal. The spectrum of $HvfBC_{fusion}$ (treated with 1.5 mol equiv. of ascorbate and desalted to rid of excess ascorbate) is shown in black, and the spectrum upon addition of HvfA (with Cys in disulfide bonds) is shown in red. *EPR conditions:* microwave frequency ~9.3 GHz, temperature 12 K, 320 ms time constant, 10 G modulation, 25 dB microwave attenuation.

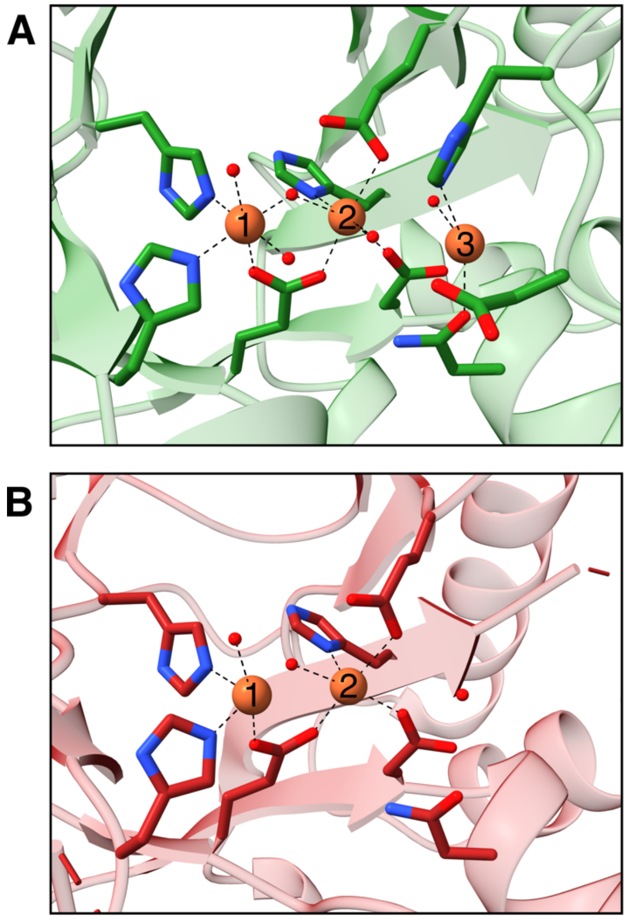

**Figure S16.** The iron-binding sites of HvfB homologs. (**A**) MbnB (PDB ID: 7TCU) with low iron occupancy (~0.26) of the third iron ion.[3] (**B**) The iron-binding site of an HvfB homolog from *Histophilus somni* 129PT (PDB ID: 3BWW).[4] Asp and His ligands to the third Fe-binding site are disordered. A buffer-derived cacodylate molecule coordinating Fe1 and Fe2 is hidden for clarity.

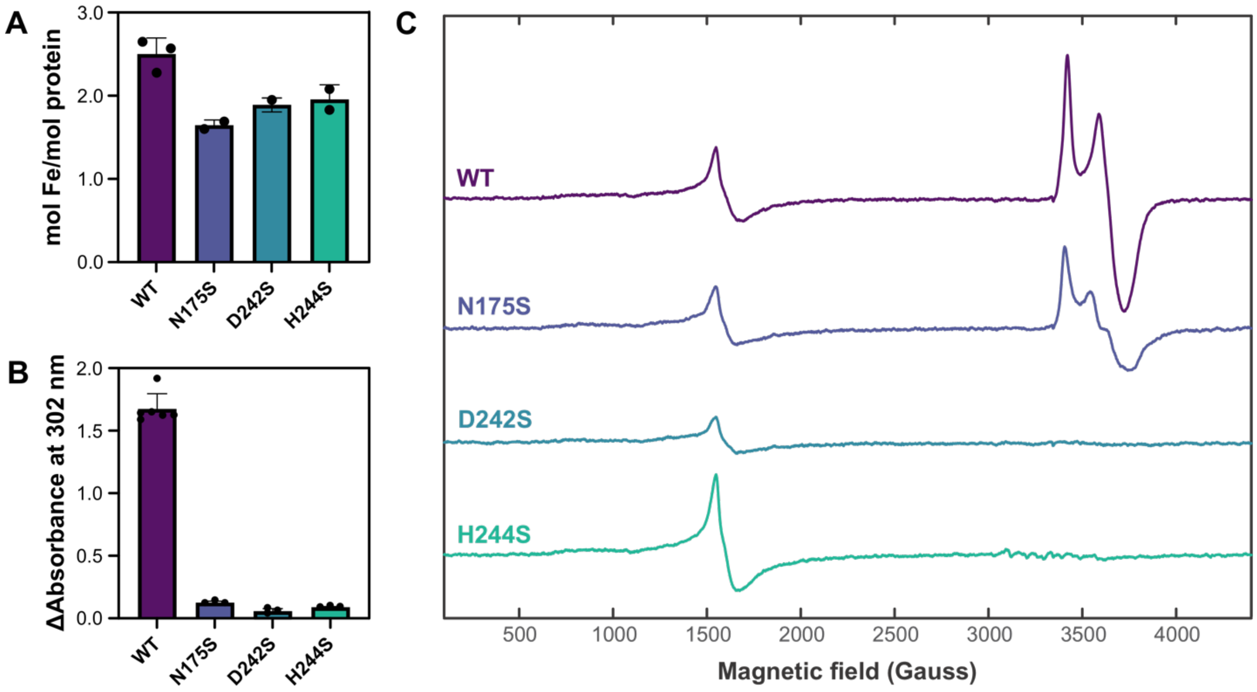

**Figure S17.** (**A**) Iron content of HvfBC$_{fusion}$ Ser variants. (**B**) Activity of HvfBC$_{fusion}$ Ser variants, measured by the increase in the product chromophore at 302 nm. Mass spectral analysis of the resulting HvfA is shown in Table S7. (**C**) X-band EPR spectra of HvfB Ser variants treated with 1 molar equivalent of sodium ascorbate. *EPR conditions:* microwave frequency ~9.3 GHz, temperature 12 K, 320 ms time constant, 10 G modulation, 25 dB microwave attenuation.

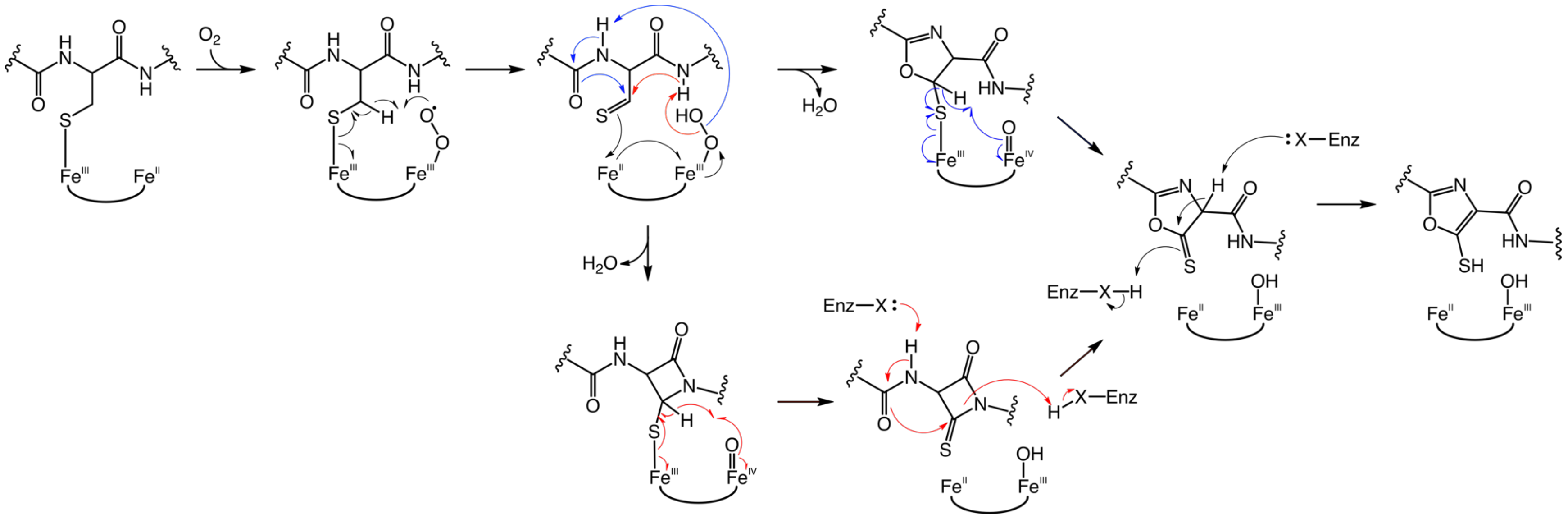

**Figure S18.** Proposed mechanism of oxygen activation and thiooxazole formation by a mixed-valent Fe(II)/Fe(III) cofactor. Substrate binding to Fe1 results in coordination by the Cys thiolate, followed by oxygen binding to Fe2 and activation to form a diferric-superoxo intermediate. This species cleaves the Cys Cβ–H bond and results in the formation of a carbon radical that combines with a thiolate electron to generate a thioaldehyde and concurrently reduce Fe1 to the ferrous state. The ensuing Fe(II)/Fe(III)-hydroperoxo can then initiate thiooxazole formation in two manners. **Red arrows:** Akin to the mechanism proposed for oxazolone formation by MbnBC,[3] the hydroperoxo pulls a proton from the *n*+1 amide, which attacks the thioaldehyde to generate a β-lactam ring. Concurrent heterolytic O–O scission and electron transfer between the iron ions result in an Fe(III)/Fe(IV)-oxo intermediate. This species abstracts another hydrogen atom from the former Cβ, and the carbon radical combines with a thiolate electron to produce a thione and regenerate the Fe(II)/Fe(III) cofactor. Enzyme base-catalyzed deprotonation of the Cys amine initiates lactam ring-opening by attack from the *n*–1 carbonyl, which forms an oxazolone-5-thione. **Blue arrows:** The Fe(II)/Fe(III)-hydroperoxo pulls a proton from the *n*–1 amide nitrogen, resulting in a 5-thiooxazoline. Heterolytic O–O cleavage and electron donation from Fe1 produces an Fe(III)/Fe(IV)-oxo species, which then abstracts a hydrogen atom from C5. The ensuing carbon radical combines with a thiolate electron to result in the oxazoline-5-thione and reform the Fe(II)/Fe(III) cofactor. Regardless of the mechanism of ring formation, it may be oxidized to the oxazole by enzyme acid/base chemistry, resulting in the final 5-thiooxazole moiety.

### References

(1) Xing, G.; Hoffart, L. M.; Diao, Y.; Prabhu, K. S.; Arner, R. J.; Reddy, C. C.; Krebs, C.; Bollinger, J. M. A coupled dinuclear iron cluster that is perturbed by substrate binding in *myo*-inositol oxygenase. *Biochemistry* **2006**, *45* (17), 5393–5401. DOI: https://doi.org/10.1021/bi0519607.

(2) Manley, O. M.; Shriver, T. J.; Xu, T.; Melendrez, I. A.; Palacios, P.; Robson, S. A.; Guo, Y.; Kelleher, N. L.; Ziarek, J. J.; Rosenzweig, A. C. A multi-iron enzyme installs copper-binding oxazolone/thioamide pairs on a nontypeable *Haemophilus influenzae* virulence factor. *Proceedings of the National Academy of Sciences* **2024**, *121* (28), e2408092121. DOI: https://doi.org/10.1073/pnas.2408092121.

(3) Park, Y. J.; Jodts, R. J.; Slater, J. W.; Reyes, R. M.; Winton, V. J.; Montaser, R. A.; Thomas, P. M.; Dowdle, W. B.; Ruiz, A.; Kelleher, N. L.; et al. A mixed-valent Fe(II)Fe(III) species converts cysteine to an oxazolone/thioamide pair in methanobactin biosynthesis. *Proceedings of the National Academy of Sciences* **2022**, *119* (13), e2123566119. DOI: https://doi.org/10.1073/pnas.2123566119.

(4) Joint Center for Structural Genomics (JCSG). Crystal structure of a duf692 family protein (hs_1138) from *Haemophilus somnus* 129pt at 2.20 A resolution. **2008**. DOI: https://doi.org/10.2210/pdb3bww/pdb.
